# Decoding the coupled decision-making of the epithelial-mesenchymal transition and metabolic reprogramming in cancer

**DOI:** 10.1101/2022.08.02.502548

**Authors:** Madeline Galbraith, Herbert Levine, José N. Onuchic, Dongya Jia

## Abstract

Cancer metastasis relies on an orchestration of multiple traits driven by different functional interacting modules including metabolism and epithelial-mesenchymal transition (EMT). Cancer cells can adjust their metabolism during metastasis by increasing oxidative phosphorylation without compromising glycolysis, acquiring a hybrid metabolic phenotype (W/O). Often required by metastasis, cancer cells engage EMT and can acquire a hybrid epithelial/mesenchymal (E/M) phenotype. Both the W/O and E/M states are associated with high metastatic potentials. Many regulatory links coupling metabolism and EMT have been identified, but how these two modules regulate each other remains largely unexplored. Here, we investigate the coupled decision-making networks of metabolism and EMT, and systematically analyzed the effect of their crosstalk. This crosstalk can exhibits synergistic or antagonistic effects on the acquisition and stability of different coupled metabolism-EMT states. Strikingly, the aggressive E/M-W/O state can be enabled and stabilized by the crosstalk irrespective of these hybrid states availability in individual metabolism or EMT modules. To acquire an E/M-W/O state, the W/O state emerges first, followed by the E/M state, suggesting metabolism can drive EMT. Our work emphasizes the mutual activation between metabolism and EMT, providing an important step towards understanding the multi-faceted nature of cancer metastasis.

## Introduction

Metastasis remains the leading cause of cancer-related deaths [1] and it is critical to understand the physiological properties of cells that migrate from the primary tumor and initiate metastatic lesions. Typically, these properties have been studied one at a time. For example, during the epithelial-mesenchymal transition (EMT) cells progressively lose epithelial (E) features such as cell-cell adhesion and apical-basal polarity, and acquire mesenchymal (M) features such as migration, invasion, and resistance to immune response [2]. The EMT has consistently been implicated in cells acquiring metastatic potential [3] and therapeutic resistance [4]. Recently, the bimodal picture of EMT has been superseded by a more complex scenario involving the hybrid epithelial/mesenchymal (E/M) phenotype which exhibits combined traits of epithelial (cell-cell adhesion) and mesenchymal (invasion) at the single-cell level. The existence of a hybrid E/M state has since been experimentally verified and associated with therapy resistance alongside poor survival rates [5–8]. Importantly, hybrid E/M cells migrate collectively and appear to be the most capable of initiating metastatic growth [8–13]. Fully understanding the behavior of the hybrid E/M phenotype is still an active area of research.

Metabolic reprogramming, another hallmark of cancer, enables cancer cells to adjust their metabolic activity for biomass and energy supply to survive in hostile environments [1,14]. Normal cells typically utilize oxidative phosphorylation (OXPHOS, O) under normoxic conditions and glycolysis under hypoxic conditions. However, cancer cells often prefer glycolysis even when oxygen is available (i.e., the Warburg effect (W) or aerobic glycolysis) [15,16]. During metastasis, cancer cells adjust their metabolic phenotype to survive in varying environments, resulting in cells switching between different types of metabolism [17– 19]. Metabolic reprogramming can enable cancer cells to combine different metabolic modes, such as acquisition of a hybrid W/O phenotype[20]. The W/O phenotype is associated with enhanced metabolic potentials, high metastatic potential [21,22], and actively uses both glycolysis and OXPHOS [23,24]. This suggests a tight connection between metabolic plasticity and cancer metastasis, specifically the hybrid W/O state with high metastatic potential.

As already mentioned, many studies of metastasis focused on either EMT or metabolism[10–13,17–19]. However, it has become increasingly clear extensive crosstalk exists between EMT and metabolism [22]. Recent studies show metabolic reprogramming can drive EMT and increase metastatic potential, or induction of EMT can drive metabolic reprogramming [25–29]. The underlying mechanisms that control how the metabolism functional module drives the EMT functional model, and vice versa, remain poorly understood, with several hypotheses discussed below. Kang et. al. suggested cancer cells typically undergo metabolic reprogramming first and then trigger EMT [30]. This coupling, presumably, is a consequence of changes in the tumor microenvironment (TME) fostering metabolic reprogramming which drives EMT [28,29]. Another hypothesis is mutual activation between EMT and metabolic reprogramming contribute to flexible coupling of various EMT states with metabolic states. Possibly, the two hybrid phenotypes (E/M and W/O) become coupled under certain crosstalk conditions, leading to a greatly increased metastatic potential [22]. Evidence supporting this connection has recently been noticed in CTCs which exhibit enhanced OXPHOS with no compromise in glycolysis [31] and they may consist mainly of hybrid E/M cells, especially at high levels of the antioxidant regulator NRF2 [32]. Additionally, hybrid E/M-like breast cancer stem cells exhibit increased levels of OXPHOS and glycolysis [33,34]. While there have been preliminary indications of the coupling of EMT and metabolic states, a systematic analysis of this coupling remains to be explored.

To decode the coupled decision-making circuit of EMT and metabolism, we developed a mathematical model which couples the core gene regulatory circuit of EMT – *µ*_34_/SNAIL/*µ*_200_/ZEB [10] with that of the metabolism one – AMPK/HIF-1/ROS [20]. By analyzing the coupled circuit, we identified the *µ*_34_/HIF-1/mtROS/*µ*_200_/SNAIL axis as a key promoter of the coupled E/M-W/O state. Additionally, HIF-1 may play a more central role in metabolism driving EMT than AMPK. Strikingly, we found the bidirectional crosstalk ensures parameter space regions exist for which only the E/M-W/O state is accessible, and the biological significance of these parameters will depend on details of the microenvironment. Interestingly, even if the individual circuits cannot give rise to the hybrid phenotype (i.e., neither the E/M or W/O states are initially accessible), upon including crosstalk, the coupled E/M-W/O state emerges. Our results therefore suggest a highly aggressive plastic phenotype along both the EMT and metabolic axes (E/M-W/O) is a likely choice for a subset of cancer cells and, speculatively, may be critical for metastasis.

### Model: Coupling the regulatory networks of EMT and metabolism

While the mechanisms of EMT and cancer metabolism have been investigated individually, the crosstalk between the two circuits and phenotypic correlations are still largely unknown. To decode the crosstalk between EMT and metabolism, we couple our previously published regulatory networks of EMT [10] and metabolism [20] by including the mutual regulatory links between these two circuits; see Fig. 1A for the coupled network and Table S5 for details of the crosstalk. The crosstalk between the EMT and metabolism circuits can be direct (e.g., HIF-1 upregulating SNAIL) or indirect (e.g., *µ*_34_ upregulating mtROS), the latter arising because our formulation focuses only on a few core components and effective interactions between them can occur via intermediate reactants. We initially focus on the core networks and investigate the role of crosstalk on the coupling of EMT and metabolism states. Then we examine whether crosstalk contributes to the emergence of the hybrid states and the stability of the E/M-W/O state.

**Figure 1.**
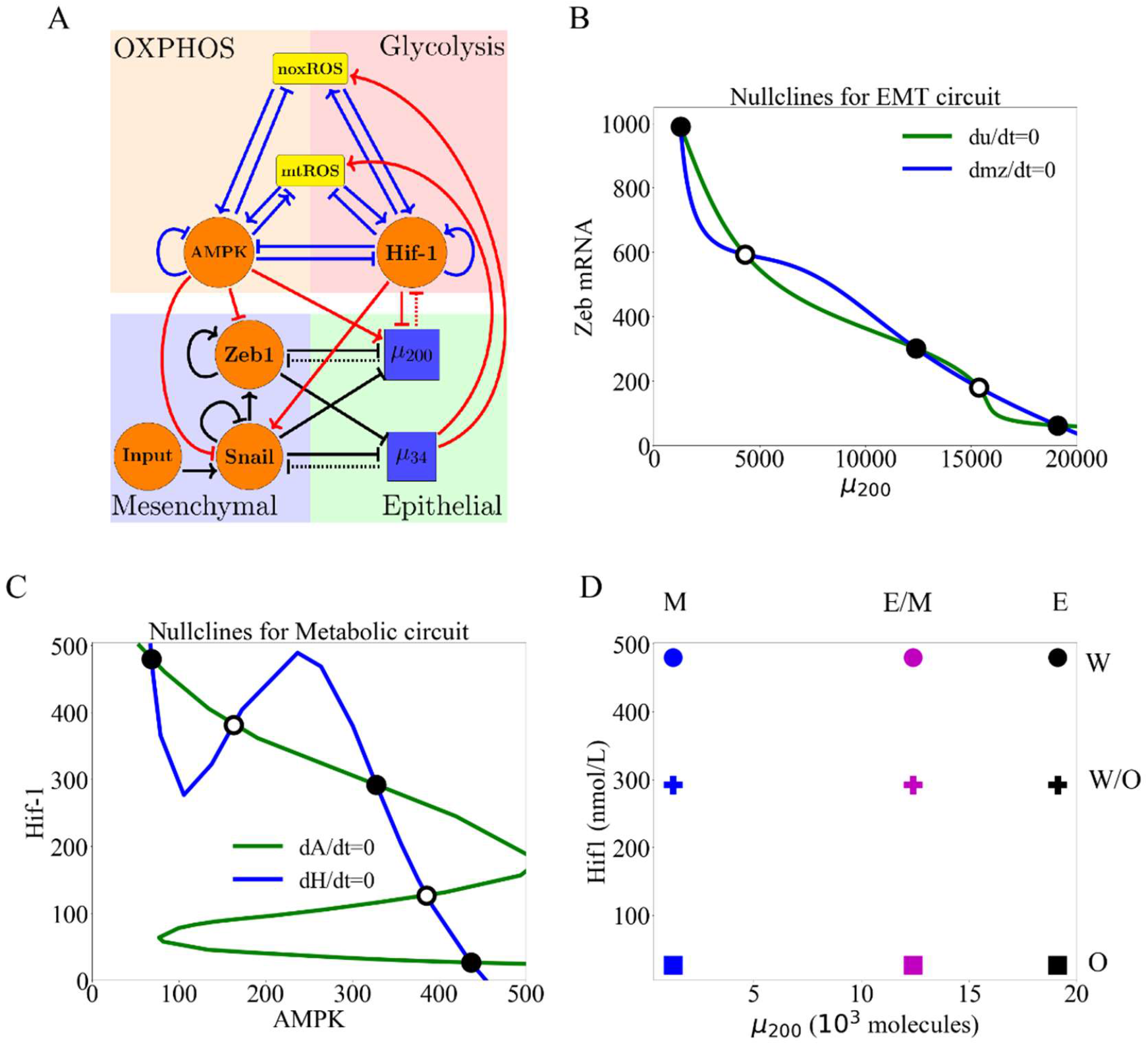
The coupled EMT/MR circuit results in nine possible steady states. **(A)** The network showing the core EMT module (bottom), the core metabolic module (top), and the crosstalk noted in red. The dashed lines denote miRNA-based regulation. Regulatory links ending in bars represent inhibition while arrows represent activation. **(B)** The nullclines of the EMT network. The system is tristable with states E (high µ_200_/low Zeb), M (low µ_200_/high ZEB), and E/M (intermediate µ_200_/ZEB). **(C)** The nullclines of the metabolic network. The system is tristable with stable states O (high AMPK/low HIF-1), W (low AMPK/high HIF-1), and W/O (intermediate AMPK/HIF-1). **(D)** The nine possible states when all crosstalks are inactive. The blue, purple, and black markers represent the EMT states. The circle, cross, and square represent the metabolism states (e.g., the coupled E/M-W/O state is represented as a purple cross).

Previous investigations presented individual insight into the core EMT and metabolism networks. Exploration of the core EMT network by Lu and collaborators showed the *µ*_200_/ZEB module was responsible for EMT tristability – epithelial (E, high *µ*_200_/low ZEB), mesenchymal (M, low *µ*_200_/low ZEB), and E/M (intermediate *µ*_200_/intermediate ZEB) – whereas the *µ*_34_/SNAIL module mainly acted as a noise buffer [10](see Fig. 1B, and section S2.1). In a separate line of investigation, a proposed regulatory circuit of metabolism AMPK/HIF-1/ROS, provided insight into cancer metabolism plasticity and switching between different metabolic phenotypes. Through this reduced circuit, Yu and collaborators show cancer cells can acquire at least three different metabolic phenotypes – OXPHOS (high AMPK/low HIF-1), Warburg (low AMPK/high HIF-1), and W/O (intermediate AMPK/HIF-1) [20] (see Fig. 1C).

To couple the regulatory circuits of EMT and metabolism, we did extensive literature search and identified the main bi-directional crosstalk between these two circuits (see Fig. 1A). Please refer to supplementary Table S5 for a detailed description of the included crosstalk. We utilize a framework consistent with the individual models of the EMT and metabolism network, treating the coupled network as a transcription-translation chimeric circuit [10]. The transcriptional regulation is mathematically represented as a shifted Hill function [35], where the fold change (*λ*) represents the magnitude of the activation (*λ* >1) or inhibition (0=<*λ* <1). For readability of the figures, we define the parameter Λ=1-*λ* such that maximal inhibition occurs when Λ=1 (*λ*=0) and no inhibition when Λ=0 (*λ*=1). We also consider the binding/unbinding dynamics (e.g., mir200 silencing HIF-1), where the functions *Y*_*μ*_, *Y*_m_, and L represent the active miRNA degradation rate, active mRNA degradation rate, and translation rate (details in SI section 1.1, Fig. S1).

The new model we propose here is built by including these crosstalk links to couple the circuits of EMT and metabolism (see SI Section 1 for the full equations, parameters and brief explanation). We started with parameters such that both the EMT and metabolic networks are tristable. When the crosstalk is inactive, at maximum nine possible combinations of the EMT and metabolic phenotypes occur: E-W, E-O, E-W/O, M-W, M-O, M-W/O, E/M-W, E/M-O, and E/M-W/O (Fig. 1D, details of numerical integration and analysis are given in section S2). By activating the regulatory links, we can identify how the crosstalk affects the coupling between EMT states and metabolism states.

First, we must develop a classification of these coupled states. While the W state is characterized by high HIF-1/low AMPK and the E state by high *µ*_200_/low ZEB expression, including the crosstalk will quantitatively alter the expression profiles for the various steady states. Therefore, the use of fixed thresholds to determine the state of the cell is no longer appropriate. Instead, we use a distance metric normalized by the expression of the decoupled network to classify the generated expression profiles as indicative of one of the nine coupled states (see Section S2.3 for details). With our baseline decoupled network parameters, we show the hybrid states (W/O and E/M) are most populous followed by the W and M states, with the O and E states being least populated (Fig. S2-S4, note the frequency of these states depends on the model parameters).

## Results

### Individual crosstalk can push the downstream circuit towards a single state

Let us start by making just one crosstalk active, e.g., caused by an EMT-related microRNA. Now, in our model there is an unaffected upstream subnetwork (EMT, from where the link originates) and a regulated downstream one. (Note the model ignores any possible dilution of the microRNA due to its action on ROS; see below).

When noxROS is upregulated by *µ*_34_ (Fig. 2A), as there is no feedback to the EMT network, the occupancy of the E, E/M, and M states are unchanged. As the level of noxROS increases, the W-associated states are lost; first the E-W state, then the E/M-W, and finally the M-W state (Fig. 2B, section S2.4). Additionally, as noxROS increases, the W/O-associated states are stabilized and little change occurs for O-associated states (Fig. 2C). Upon analyzing the coupled states, if noxROS increases then the E/M state becomes more likely to be associated with the W/O state (Fig. 2D). Additionally, the W state is most associated with the M state which is expected since the M state has the lowest *µ*_34_ level, resulting in a smaller increase of noxROS compared with other EMT states (Fig. 2E).

**Figure 2.**
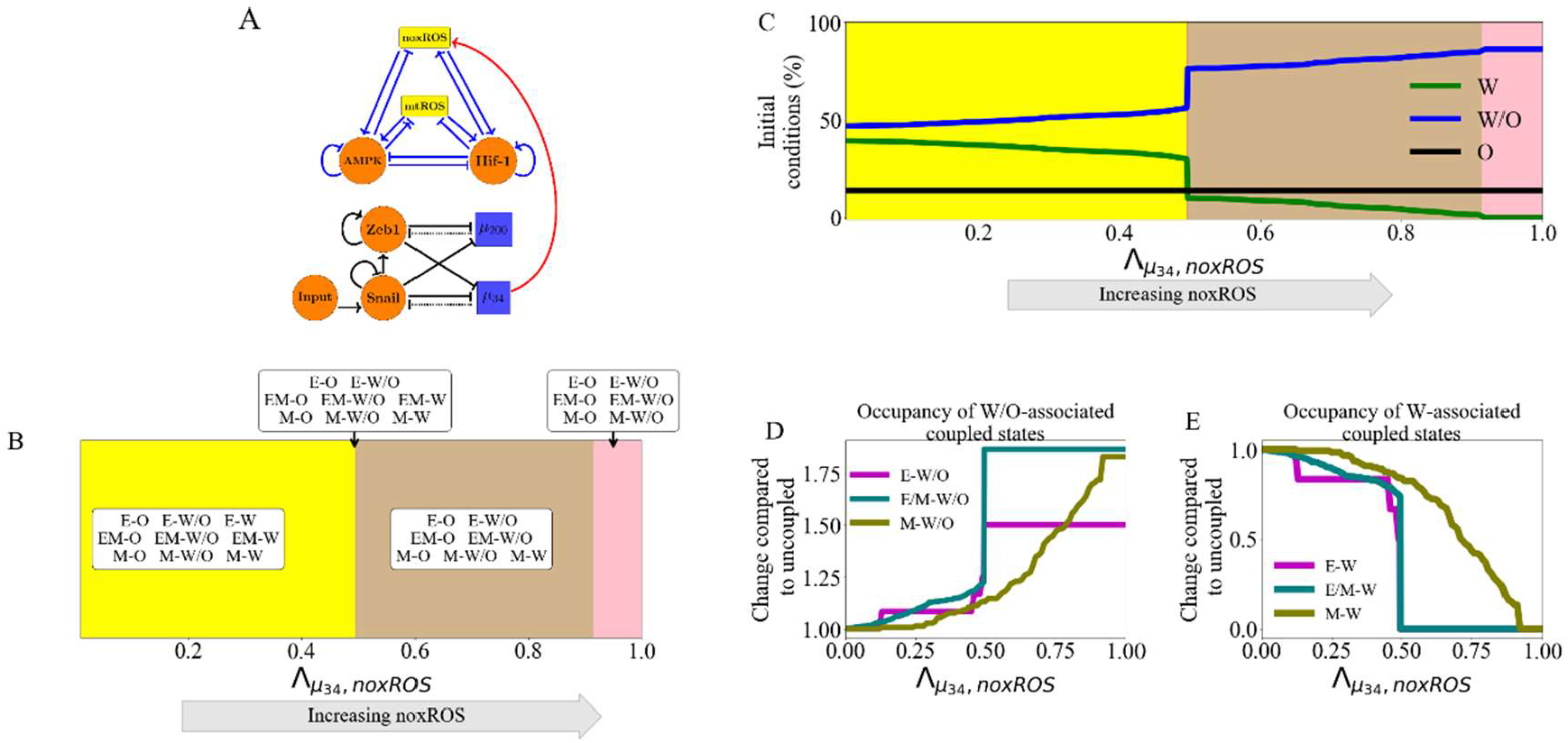
noxROS upregulated by μ_34_ stabilizes the W/O state and enhances the E/M-W/O coupled state. **(A)** A diagram of the core EMT (bottom) and metabolic (top) circuits connected by the crosstalk *µ*_34_ upregulating noxROS (red link). The EMT network is unchanged, as there is no feedback. **(B)** As *µ*_34_ upregulates noxROS, 4 distinct phases occur; all nine couple states followed by loss of the E-W, E/M-W, and M-W states (yellow, red, tan, and pink regions, respectively). **(C)** The lines represent the number of initial conditions leading to the W (green), O (black), or W/O (blue) states as noxROS increases. (Background colors correspond to the phase colors of (B).) **(D)** The frequency of the W/O-associated states (i.e., E-W/O, M-W/O, and E/M-W/O) compared to the inactive system 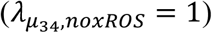. All W/O-associated states are promoted, with the E/M-W/O state being greatly increased once 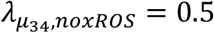. **(E)** Same as (D) but for W-associated states. The W-associated states are suppressed with the M-W state lasting longest.

Similar changes have been observed via *µ*_34_ upregulating mtROS. Consistent with upregulating noxROS, the E-W and E/M-W states are suppressed first and the E/M-W/O state is stabilized (Fig. S5). Additionally, upregulating mtROS exhibited a greater increase in the E/M-W/O state than upregulating noxROS. Further, activation of mtROS stabilizes the W/O state but reduces occupancy of the O state and W state. Together, these results suggest mtROS and noxROS may be critical factors in regulating the coupling of two hybrid states (E/M-W/O).

### Regulation of HIF-1 affects both subcircuits

While the *µ*_34_ links only affect the downstream network, the miRNA regulation of HIF-1 by *µ*_200_ can affect both networks. This arises in our model because *µ*_200_ mediates the transcription and translation of HIF-1 mRNA, and as a result, *µ*_200_ can be recycled or degraded. Therefore, while the downstream metabolic network is modulated, the upstream EMT network is also affected via change of *µ*_200_. We have defined a silencing function *P*_*H*_(*µ*) to simulate the above-mentioned effect of *µ*_200_ on HIF-1 (details of function in section S2.5). Note as silencing increases, the restriction of the EMT states occur immediately; close to *P*_*H*_(*µ*) = 0, the only EMT state allowed is M while all the metabolic phenotypes are allowed. As *µ*_200_ silences HIF-1, the W/O and W states are suppressed sequentially, and the O state is promoted. Additionally, as HIF-1 levels decrease, less degradation of *µ*_200_ (caused by binding to HIF-1 RNA) occurs, resulting in gradual disappearance of the M state. Thus, when HIF-1 mRNA is fully silenced, only the E-O and E/M-O states remain (Fig. S6). Since the E/M state does not reappear until after the metabolic system has fully transitioned to O, the E/M-W/O state is not observed for any value of *µ*_200_ silencing HIF-1 mRNA. These results suggest *µ*_200_ overexpression could promote the O-associated states (E-O and E/M-O) and destabilize the coupled E/M-W/O state.

### Inclusion of multiple miRNAs of the EMT network can stabilize the hybrid W/O phenotype

We next wish to determine how including links emanating from both *µ*_200_ and *µ*_34_ can synergistically drive metabolic reprogramming and promote the coupled E/M-W/O state. While upregulating ROS causes an increase in the E/M-W/O state, we showed *µ*_200_ silencing HIF-1 suppresses the E/M-W/O state; therefore, we expect some suppression of the E/M-W/O state when including both μ_200_ and μ_34_ crosstalks. Interestingly, the E/M-W/O state can be fully suppressed when decreasing HIF-1 and upregulating noxROS, but only partially suppressed when decreasing HIF-1 and upregulating mtROS (Fig. S7). These results suggest the type of ROS present has different effects on the existence of the E/M-W/O state.

The E/M-W/O state is stabilized if mtROS is upregulated, but upregulating noxROS has minimal effect on the E/M-W/O state. Strikingly, when all miRNA crosstalks are active (*µ*_200_ silencing HIF-1, and *µ*_34_ upregulating noxROS and mtROS, Fig. 3A) the E/M-W/O state can be suppressed even if the W/O state is present (Fig. 3B). Further, the E/M-W/O state is present for all values of noxROS but is only present at increased levels of mtROS (Fig. 3B-C and S8).Additionally, at high levels of mtROS, the E/M and W/O states are likely to be associated, suggesting upregulating mtROS can stabilize the E/M-W/O state (Fig. 3C). Depending on the initial conditions, the E/M-W/O state is accessible if HIF-1 is partially silenced and at high levels of noxROS and mtROS (Fig. 3D). This suggests there is a synergistic effect between the crosstalks, resulting in an increased parameter space enabling the E/M-W/O state than expected based on the individual crosstalks. Further, the difference in the effect of noxROS and mtROS seems to result from the frustrated regulation of mtROS by HIF-1 and *µ*_34_. Therefore, feedback loops between mtROS, HIF-1, *µ*_34_, and *µ*_200_ together control the appearance of the E/M-W/O state.

**Figure 3.**
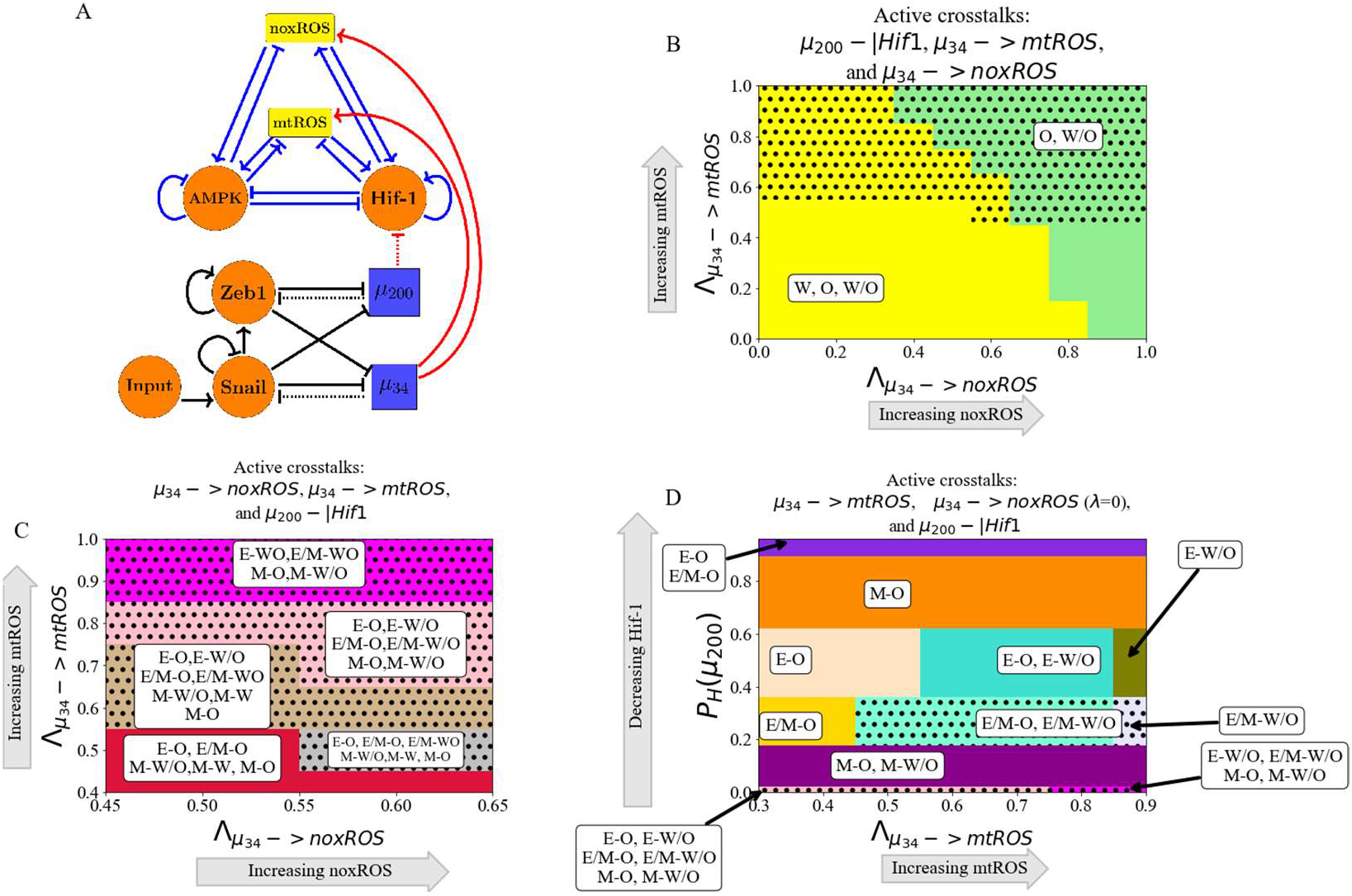
μ_200_ and μ_34_ can upregulate the W/O phenotype. The E/M-W/O state can also be upregulated and mtROS seems to be a key regulator.**(A)** Schematic illustration of the coupled metabolic (top) and EMT (bottom) networks with all miRNA-mediated regulatory links active (*µ*_34_ upregulating mtROS, *µ*_34_ upregulating noxROS, and *µ*_200_ silencing HIF-1). **(B)** The phase plane corresponding to all miRNA-mediated links (pictured in A). In this phase plane, *µ*_200_ silencing HIF-1 corresponds to the rightmost blue region of Fig. S6 (all metabolic phenotypes are possible), increased noxROS suppresses the W state, and increasing mtROS causes the E/M-W/O coupled state to appear (black dotted region). **(C)** The coupled states of (B), zoomed in on the middle region. The E/M-W/O state exists when mtROS is upregulated. **(D)** At maximum noxROS levels, increased mtROS levels (x-axis), and moderately silenced HIF-1 (y-axis) there are regions where the E/M-W/O state is possible (black dotted regions).

### Metabolic reprogramming can drive EMT

We next consider information flowing in the opposite direction, from metabolism to EMT and determine the effect of each metabolism-driven crosstalk on the coupled states. First, we analyzed the links in which HIF-1 upregulates SNAIL (Fig. 4A and S9) or inhibits *µ*_200_ (Fig. S10). As expected, both HIF-1 mediated links push the system towards the M state. Further, both the E and E/M states are most associated with the O state (when the HIF-1 level is relatively low) while the M state is initially associated with the W state. This correlation between the E-O and M-W states is assumed in much of the literature [24]. Similarly, modulating the EMT-inducing signals, such as TGF-β, that activate SNAIL can alter the stability of the E/M state and the coupled states (see Fig. S11). Opposite to the HIF1-mediated crosstalks, AMPK-mediated crosstalks (upregulating *µ*_200_, downregulating SNAIL, or downregulating ZEB) push the EMT network to adopt an E state and suppress the E/M state, followed by the suppression of the M state (Fig. 4B and S12-S14). Additionally, when AMPK regulates the EMT circuit alone, the E and M states are still most associated with the O and W states, however, the E/M state is associated with the W state. This is in direct contrast to HIF-1 driven crosstalk in which the E/M state is coupled with O state. The results suggest the E/M state has metabolic plasticity because neither OXPHOS nor Warburg metabolism automatically associate with it.

**Figure 4.**
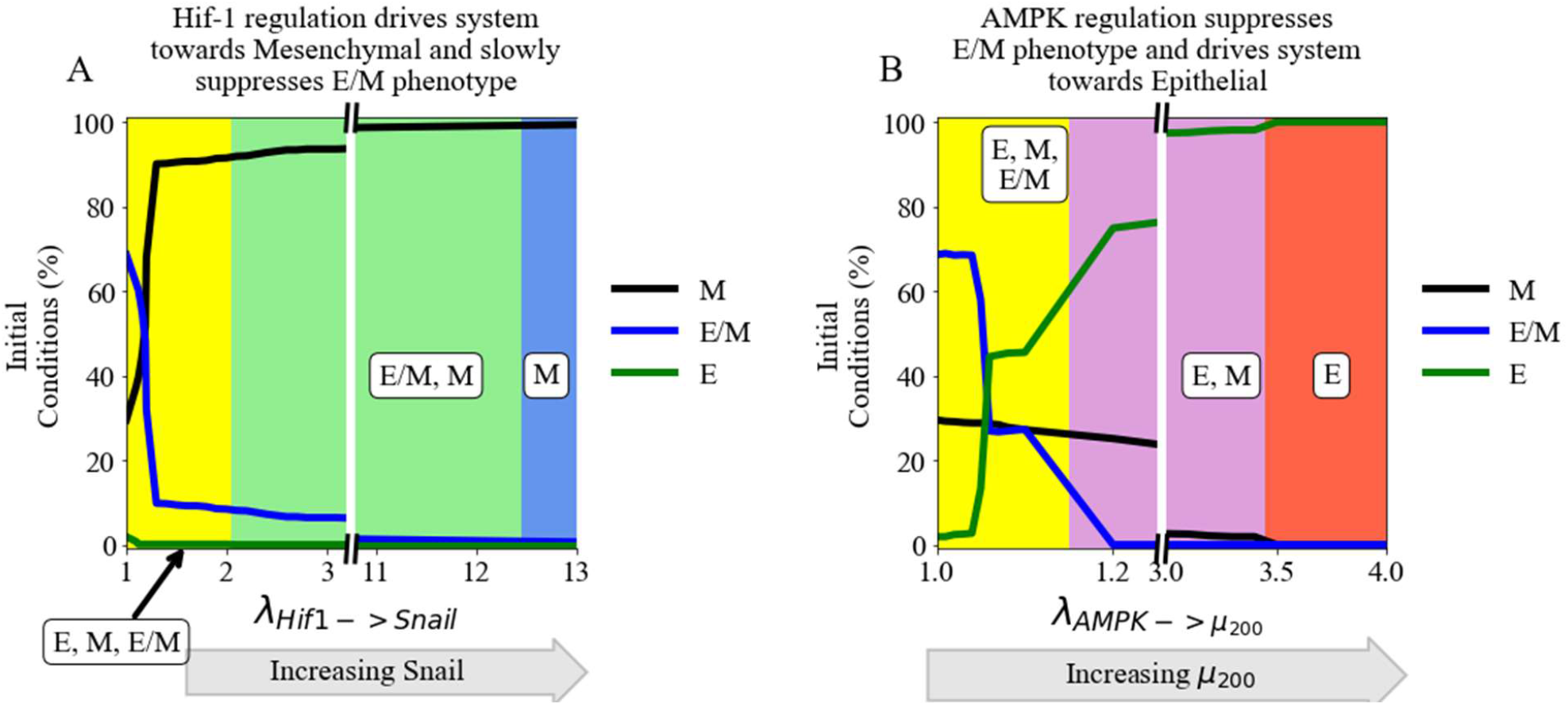
The role of metabolism in driving EMT. HIF-1 mediated crosstalks drive the EMT circuit towards the M state, while AMPK mediated crosstalks drive the EMT network towards the E state. **(A)** The frequency of the E/M, M, or E state as HIF-1 upregulates SNAIL and drives the EMT network towards mesenchymal. **(B)** The frequency of the E/M, M, or E state as AMPK upregulates *µ*_200_ and drives the system towards epithelial. The E/M state exists for larger portions of the parameter spaces for HIF-1 regulation than for AMPK-mediated crosstalks.

### TFs of the metabolic network can stabilize the E/M phenotype

Two distinct events are at play when the metabolic network regulates the EMT circuit. AMPK regulation quickly suppresses the E/M state and pushes the system towards the E state, whereas HIF-1 regulation can allow the system to maintain the E/M state for a range of strengths while ultimately pushing the system towards the M state (Fig. 4A and 4B). Thus, HIF-1 and AMPK-mediated crosstalk should act antagonistically to stabilize the hybrid state.

When at least one of the AMPK crosstalks and one of the HIF-1 crosstalks are activated, the E/M state is stabilized. Additionally, if AMPK and HIF-1 target different EMT-TFs, the E/M-W/O state may exist in larger parameter spaces than if they target the same EMT-TF (Fig. S15). This suggests activating multiple crosstalks and targeting multiple TFs is likely to stabilize the E/M-W/O state. Therefore, if all HIF-1 and AMPK-mediated crosstalks are active (Fig. 5A) then significant regions occur in which the E/M state exists (Fig. 5B). However, the E/M-W/O state only exists in a small region where *µ*_200_ is minimally upregulated. Moreover, HIF-1 driven crosstalks can maintain the E/M state longer than AMPK driven crosstalks suggesting, the reduction of the E/M-W/O state is likely due to the suppression of the E/M state by AMPK-mediated crosstalks, as mentioned above (see Fig. S12-S14). This suggests HIF-1 driven crosstalk is more strongly correlated with the E/M state than AMPK driven crosstalk.

**Figure 5.**
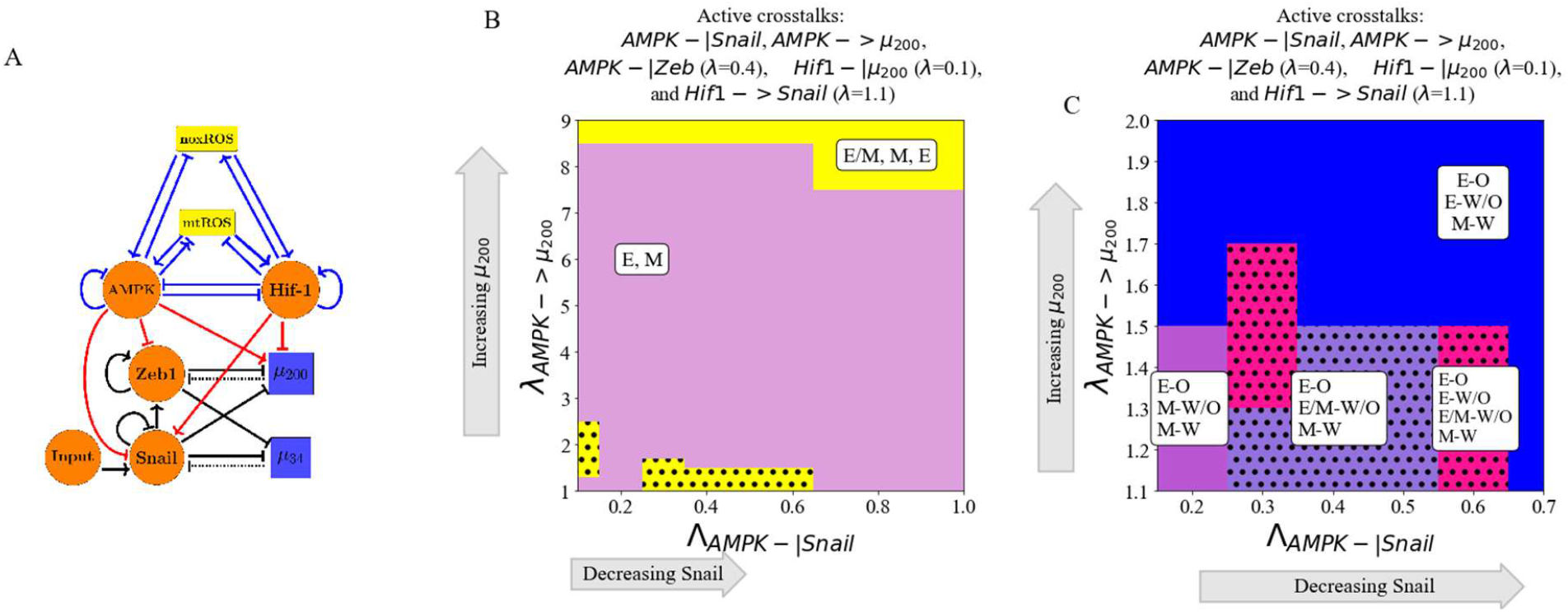
AMPK and HIF-1 cooperate to upregulate the hybrid E/M state. When all HIF-1 and AMPK controlled crosstalks are active (HIF1->Snail, HIF1-|*µ*_200_, AMPK-|Snail, AMPK-|Zeb, AMPK-> *µ*_200_) the E/M-W/O state can be stabilized. The metabolism-mediated crosstalks work antagonistically to stabilize the E/M state. **(A)** Schematical illustration of the network showing the metabolism-mediated crosstalk. **(B)** The phase plane of potential EMT states when all metabolic driven crosstalks are active. The E/M state is only accessible when *λ*_AMPK->_ μ_200_ is near 1 or very high (i.e., at the extremes of regulation). **(C)** The coupled states when the EMT circuit is regulated by the metabolic circuit (pictured in A). The results suggest a direct correlation between the E, E/M, and M states to the O, W/O, and W states, respectively.

If all EMT regulating crosstalks are active, then there are regions where the E/M-W/O state exists. Additionally, the E state is typically coupled to the O state (E-O), the M state is associated with the W state (M-W), and when the E/M state is present it is typically associated with the hybrid W/O state (Fig. 5C). In fact, for any system, if only three coupled states are available and each has a distinct phenotype of the EMT and metabolic networks, then the only possible set of states is E-O, M-W, and E/M-W/O. This behavior represents the full coordination of EMT and metabolism and suggests clusters of migrating cells utilize a combination of aerobic glycolysis and OXPHOS. Given tumors are metabolically heterogeneous, this result suggests the topology and parameters of the system may only represent certain microenvironments and is a limitation of our study.

### The Hybrid E/M-W/O phenotype

Recently, it has been suggested the most aggressive cancer phenotype is characterized by the hybrid E/M or W/O states [24]. Therefore, we now focus on how the crosstalk between EMT and metabolism networks affects the E/M-W/O state.

The hybrid E/M-W/O state can be promoted for multiple combinations of crosstalk. For example, the E/M-W/O state can be stabilized when AMPK downregulates SNAIL, HIF-1 downregulates *µ*_200_, and *µ*_34_ upregulates mtROS (Fig. 6A and S16A). Further, the E/M-W/O state become more prevalent when replacing HIF-1 downregulating *µ*_200_ by increasing the EMT inducing signal to SNAIL (Fig. 6B and S16B). Interestingly, while the E/M-W/O state was stabilized in both cases (Fig. 6A-B), neither set of crosstalks could enable only the E/M-W/O state. However, it is possible to enable only the E/M-W/O state with just three regulatory links; HIF-1 inhibiting *µ*_200_, *µ*_34_ upregulating mtROS, and modulating the EMT-inducing signal (Fig. 6C and S16C). Additionally, this region, which only includes the E/M-W/O state, persists if all crosstalks are activated (Fig. 6D and S16D).

**Figure 6.**
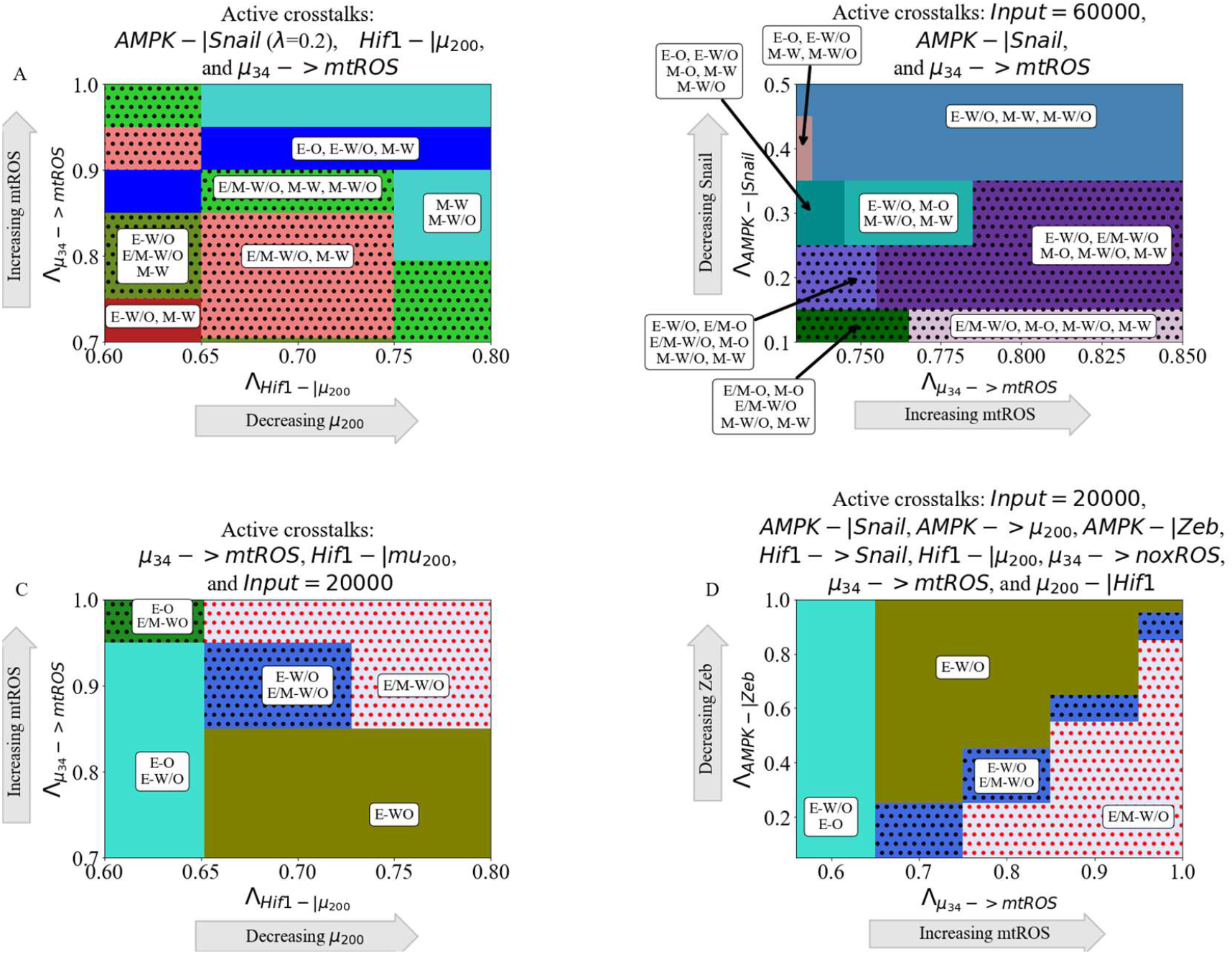
The coupling of the EMT and metabolic regulatory networks can enable a coupled hybrid E/M-W/O state. Minimally, three links (one effecting the metabolic network and two controlling the EMT network) are necessary to enable only the E/M-W/O state. **(A)** Phase diagrams of the coupled states when considering three crosstalks; the Input=60000 molecules, AMPK downregulates SNAIL, and *µ*_34_ upregulates mtROS. The E/M-W/O state is promoted when mtROS levels are increased. **(B)** The phase diagram of the coupled states when considering AMPK inhibiting SNAIL, HIF-1 inhibiting μ_200_, and *µ*_34_ upregulating mtROS. The E/M-W/O state is stabilized for some regions. **(C)** When considering the bi-directional regulation between EMT and metabolism by the three minimally necessary regulatory links (*µ*_34_ upregulating mtROS, HIF-1 inhibiting *µ*_200_, and an EMT-inducing signal on SNAIL) parameter regions exist enabling only the E/M-W/O state. **(D)** When all crosstalks are active there are regions where only the E/M-W/O state exists. Similar sets of phases in (C) and (D) suggest a progression drives the system towards the E/M-W/O state.

The proximal phases of the phase enabling only the E/M-W/O state suggests stabilization of the E/M-W/O state requires mutual activation between metabolism and EMT. When the E/M-W/O state is the only available coupled state, the surrounding phases (E-O and E-W/O) are the same whether only three crosstalks (Fig. 6C) or all crosstalks (Fig. 6D) are active, suggesting there may be a sequential path to generate the E/M-W/O state. Further, if the E/M-W/O state is not the only allowed state (Fig. 6A-B), the surrounding phases include both E-associated and M-associated states. Together, the results suggest to reach the E/M-W/O state, epithelial cancer cells first undergo metabolic reprogramming (acquiring the E-W/O state), followed by partial EMT (E/M-W/O). Although it is outside the scope of this manuscript, other crosstalk combinations may also stabilized the E/M-W/O state, and based on these results we would expect HIF-1 suppressing *µ*_200_ and *µ*_34_ upregulating mtROS to be prominent among all such combinations.

### Hybrid phenotypes are enabled by crosstalk in cells initially without the E/M or W/O state

To investigate whether the crosstalk between EMT and metabolism promotes cancer plasticity (e.g., by acquiring the hybrid states) we simulate scenarios where the individual EMT and metabolism networks cannot acquire a hybrid state. This scenario corresponds to normal physiological conditions where we expect most cells will be restricted to a binary choice of E versus M and W versus O [36] (see Fig. S17). Then we systematically analyze whether any crosstalk can enable the hybrid state to emerge.

To analyze how the EMT network can drive metabolic reprogramming, we first kept the individual metabolic circuit as a bistable system where only the W and O states are available (i.e., no W/O state). Through gradually activating the miR-34-mediated links, we found the hybrid W/O state emerges and the E/M-W/O state is stabilized when *µ*_34_ upregulates mtROS (Fig. 7A) but they do not appear when upregulating noxROS or downregulating HIF-1 (Fig. S18). This suggests, noxROS may play a context-dependent role on the coupled state, while mtROS often stabilizes the E/M-W/O state.

**Figure 7.**
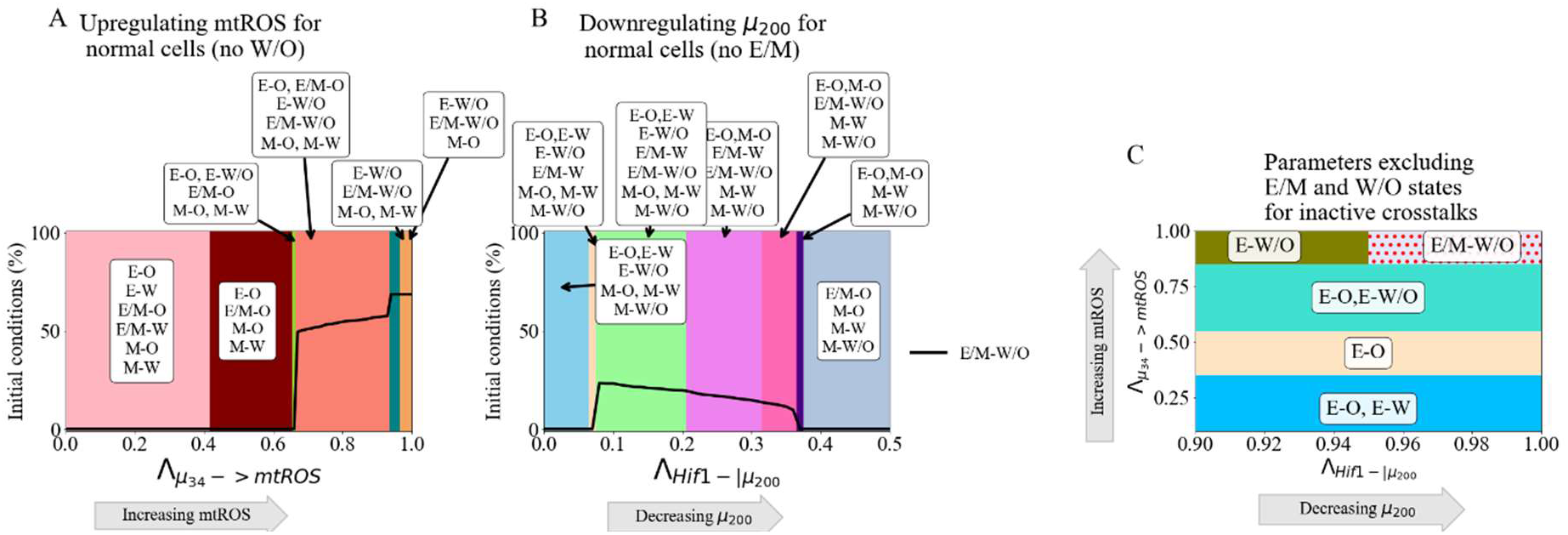
Crosstalk can generate the hybrid states. The activation of a single crosstalk can generate the hybrid state of the downstream network (W/O or E/M). **(A)** The phase diagram showing coupled states for the bistable metabolism network (O or W when the crosstalk is inactive 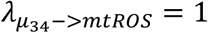.). Once mtROS is increased (near *λ*_μ3_ = 0.35), there is a sharp change with the hybrid W/O state becoming the most often occupied state. **(B)** The phase diagram of coupled states when the hybrid E/M state is not available initially when the crosstalk is inactive. As mir200 decreases, the E/M state becomes accessible. **(C)** Combining the models from (A) and (B), we generate a network which only has 4 possible coupled states if the crosstalk is inactive (E-O, E-W, M-O, and M-W). At maximum upregulation of mtROS and downregulation of *µ*_200_, only the E/M-W/O state is enabled, similar to Fig. 6C.

Next to see how metabolic reprogramming can possibly drive the hybrid E/M state, we set the EMT network to be bistable (i.e., unable to acquire the E/M state). We find the E/M state can be generate and coupled with the W/O state when HIF-1 inhibits *µ*_200_ (Fig. 7B) or HIF-1 upregulates SNAIL (Fig. S19). This suggests the master regulator of glycolysis, HIF-1, can drive cells towards the hybrid E/M state. Conversely, an individual AMPK-mediated crosstalk is unable to generate the hybrid E/M state (Fig. S19). Additionally, as with the tristable networks, two competing crosstalks (e.g., AMPK upregulating SNAIL and HIF-1 downregulating *µ*_200_) can stabilize the hybrid E/M-W/O (Fig. S20).

When both networks are in the parameter regime where the hybrid state is not available (i.e., neither the E/M or W/O state), the crosstalk can still enable the emergence of these hybrid states. Recall for the coupled tristable circuits, the simplest set of crosstalk with a parameter region enabling only the E/M-W/O state consisted of three regulatory links; HIF-1 inhibiting *µ*_200_, *µ*_34_ upregulating mtROS, and EMT-inducing signaling acting on SNAIL. When these same links are active for the bistable EMT and metabolism circuits, there are parameter spaces where only the E/M-W/O state are enabled, the results qualitatively agree with the tristable circuit results (Fig. 7C and S21 compared to Fig. 6C).

In summary *µ*_34_ upregulating mtROS can generate the W/O state and upregulate the E/M-W/O state when the metabolism circuit itself can only acquire the W and O states.Conversely, a HIF-1 mediated crosstalk can generate the E/M state and stabilize the E/M-W/O state even when the EMT circuit itself cannot acquire the E/M state. If both networks are bistable it is possible to enable only the E/M-W/O state with three regulations. These results suggest the E/M-W/O state can be generated and promoted by the crosstalk, independent of the initially available states.

### GRHL2 and OVOL can stabilize the coupled E/M-W/O state

Previously, we reported transcription factors, such as OVOL and GRHL2 can stabilize the hybrid E/M state [7,37], referred to as the phenotypic stability factors (PSFs) of the E/M state. We are curious how these PSFs regulate the coupling of the E/M state with metabolism states, specifically the E/M-W/O state. We extended the original coupled EMT-metabolism network by including OVOL and GRHL2 (Fig. 8A, parameters and modified equations of the PSF stabilized network are in Section S1.6). When a single crosstalk is active, the E/M-W/O state can exist for the entire parameter space when one of the following crosstalk is active - AMPK downregulating SNAIL, AMPK upregulating *µ*_200_, *µ*_34_ upregulating mtROS, or *µ*_34_ upregulating noxROS (see Fig. S22). The E/M-W/O state is also stabilized when HIF-1 downregulates *µ*_200_ or upregulates SNAIL (Fig. S22).

**Figure 8.**
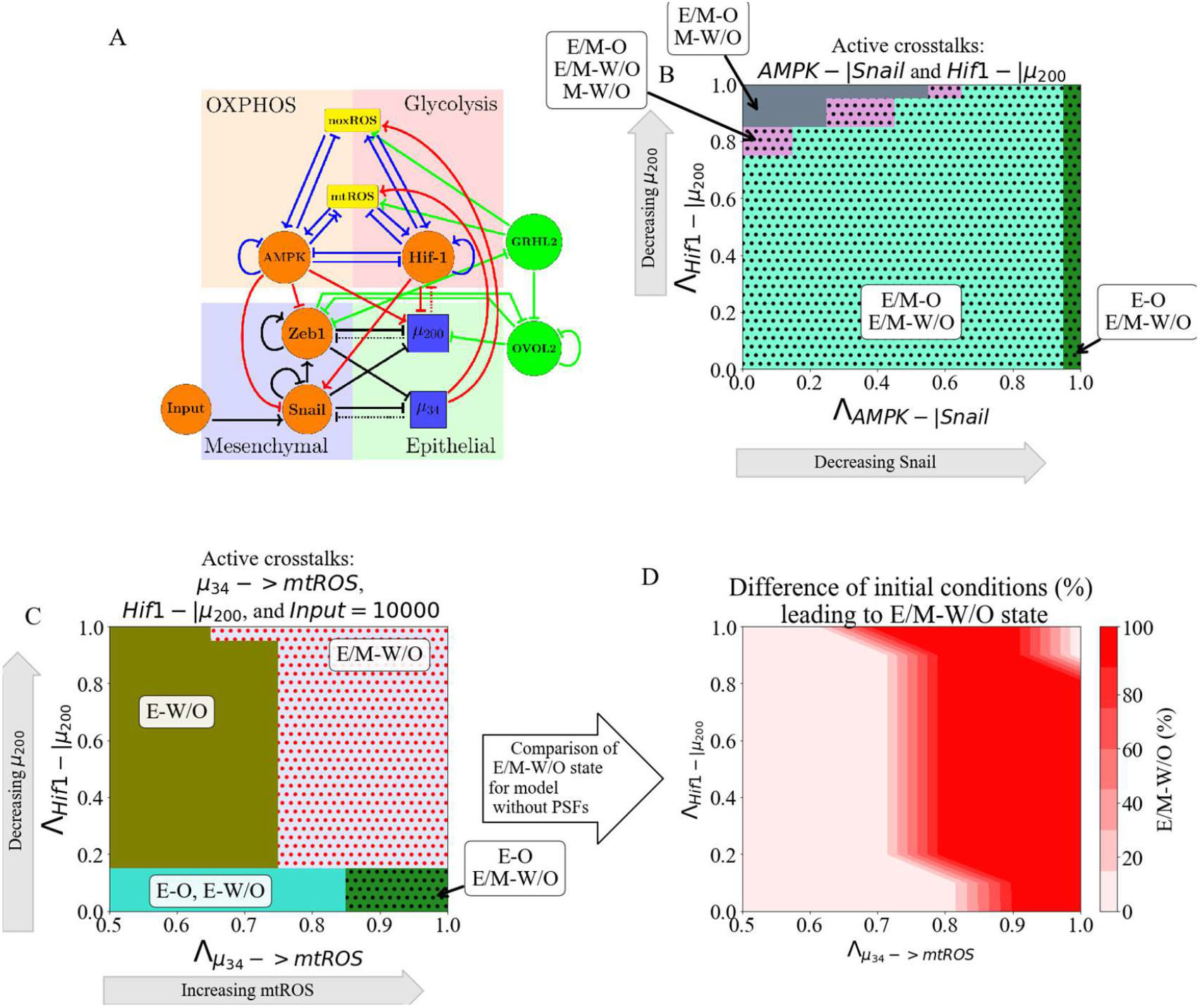
PSFs stabilizing the E/M state can stabilize the association of the E/M state with the W/O state. **(A)** The coupled EMT-metabolism network including PSFs GRHL2 and OVOL2. **(B)** The phase diagram of the coupled states shows the E/M-W/O state is present in a larger parameter space due to the PSFs stabilizing the E/M state. **(C)** The phase diagram of the coupling states when the three links (*µ*_34_-> mtROS, HIF1-|*µ*_200_, and reducing the EMT-inducing signal) are active. The parameter regime of the phase enabling only the E/M-W/O state is significantly enlarged compared to the coupled network without PSFs. **(D)** The difference in the frequency of the E/M-W/O state between (C, PSFs present) and the original model (Fig. S23, PSFs not present). The dark red region shows the phases where only the PSF stabilized model can enable the E/M-W/O state, whereas the top right corner is the region where only the E/M-W/O state enabled irrespective of the presence of PSFs.

Next, we studied the effect of the PSFs when multiple crosstalks are active. If two competing crosstalks acting on the EMT circuit are active (e.g., one HIF-1 and one AMPK mediated crosstalk), the E/M-W/O state is available for most of the parameter space (Fig. 8B). Further, the E/M-W/O state can be stabilized by activating the three crosstalks (i.e., *µ*_34_ upregulating mtROS, HIF-1 downregulating *µ*_200_, and including the EMT-inducing signaling acting on Snail) that enable only the E/M-W/O state in the absence of these PSFs. Additionally, the parameter space enabling only E/M-W/O is much larger in the presence of the PSFs (Fig. 8C-D and S23) relative to the absence of the PSFs. Further, the coupled states in the phases surrounding the phase containing only the E/M-W/O state are the same as those identified in the absence of PSFs (Fig. 8B). These results show the PSFs can stabilize the coupled E/M-W/O state.

## Discussion

Cancer malignancy relies on the orchestration of multiple hallmarks driven by different functional modules, such as metabolism and stemness [1]. It has become increasingly clear different hallmarks of cancer are extensively coupled. In this work, we focused on how reprogrammed cancer metabolism is coordinated with cancer metastasis. As EMT is often employed by cancer as part of the metastatic process, we analyzed the mutual regulation between metabolism and EMT through coupling their corresponding gene regulatory circuits. We systematically analyzed the effect of both individual and multiple crosstalk on each of the nine coupled states. The stability of the coupled states was found to vary depending on which crosstalk was active, and multiple crosstalks could exhibit synergistic or antagonistic effects.

Therefore, we primarily focused on the E/M-W/O state, as we expect these cells to be the most metastatically capable. We found (1) the E/M-W/O state can be stabilized by a single crosstalk mediated by miR-34 or two antagonistic EMT-driven crosstalks; (2) the similarities between the effects of different crosstalk (e.g., HIF1 suppressing μ_200_ compared to HIF-1 upregulating SNAIL) suggest a degree of consistency in how EMT drives metabolic reprogramming, and vice versa; (3) if crosstalk is bidirectional, it is possible to enable only the E/M-W/O state and this stabilization can be facilitated even under conditions when the individual core circuits do not generate hybrid states; (4) the E/M stabilizing PSFs (OVOL, GRHL2) also stabilize the coupled E/M-W/O state. Together, the results highlight the vital role of the EMT-metabolism crosstalk in mediating cancer metastasis.

The results of our model suggest metabolic reprogramming can drive EMT, but metabolic reprogramming does not have to be complete before EMT begins; this feature allows stabilizing of the most aggressive E/M-W/O state. Further, we identified a scenario wherein the system can follow a progression from the E-O state, first undergoing metabolic reprogramming while maintaining epithelial characteristics (E-W/O state), before undergoing partial EMT to stabilize the E/M-W/O state. Strikingly, the prevalence of the E/M-W/O state is increased by EMT-metabolism crosstalk regardless of initial phenotypic availability (i.e., whether the initial system is significantly E/M-W/O or only E-O, E-W, M-O, and M-W). Therefore, our current model provides a possible explanation for the mutual activation of metabolic reprogramming and EMT, depending on the initiating signal.

Our findings indicate all else being equal, undergoing EMT tends to correlate with using additional glycolysis. This qualitatively agrees with a recent pan-cancer study based on NCBI GEO microarray datasets and other studies [24,38]. We find HIF-1 (a marker of glycolysis) is strongly associated with EMT, suggesting the E/M state can be stabilized if HIF-1 (glycolysis) is upregulated. Additionally, our model predicts the coupling of the hybrid E/M state and high glycolysis/high OXPHOS (W/O). Notably, our model is unable to explain the cases wherein low glycolysis metabolism is correlated with EMT. However, extending the model to explicitly include coupling with additional metabolic pathways[39] may be able to explain the low glycolysis states of the pan-cancer study [38].

The coupling of the E/M and W/O states is somewhat surprising given the widespread impression that primary tumors often exhibit the Warburg effect, possibly because of their need to limit the amount of ATP produced in favor of maximizing biomass production and growth (see [40] and references therein). However, this finding is consistent with the general idea that moving from E to E/M correlates to increasing stemness, and stem-like capabilities often rely on glycolysis. It is also consistent with HIF-1 activation diminishing OXPHOS while driving EMT. Note this tendency might be over-ridden for cells requiring sufficient energy production to enable motility, such as leader cells. One possibility is that during the transition between E-O to E-W/O, when cells first become malignant, the Warburg effect is activated. Then as cells undergo EMT they tend to switch to more W until reaching a mesenchymal-like E/M state with mostly W. Lastly, as cells complete EMT and fully differentiate, they revert back to using mostly OXPHOS. The connection between EMT and metabolism may also depend on other external signals, such as the level of oxygen in the TME. For example, mesenchymal cells that reduce proliferation and have to traverse the extracellular matrix should switch to more OXPHOS, whereas ones that become quiescent in a hypoxic metastatic niche should favor glycolysis. Resolution of this issue must await a more precise idea of the phrase ‘all else being equal’.

The importance of the *µ*_34_/*µ*_200_/HIF-1/ROS/SNAIL axis for the regulation of the E/M-W/O state arises from our analysis. Our results suggest mtROS is critical for the metabolic activation of EMT, in agreement with recent experimental work that posited mtROS can drive EMT [41], control cancer invasiveness [42], and have a stronger role than noxROS [41,43]. Our results also suggest the mtROS/HIF-1 axis is critical to stabilizing the highly aggressive E/M-W/O state, and this axis has previously been associated with hypoxia-induced cancer aggressiveness [44]. Additionally, both mtROS and HIF-1 are controlled by the miRNAs of the EMT network, μ_34_ and μ_200_, confirming the importance of miRNAs in mediating the coupling of EMT and metabolism [45]. While we have parametrized the model with values from literature whenever available to ensure biological relevance, one limitation of this study is knowing how these results translate to experimental cancer studies. Thus, the significance of the mtROS/HIF-1 feedback loop should be experimentally tested by e.g., modulating ROS level via antioxidant factors such as NRF2, or modulating hypoxia to perturb HIF-1.

In line with the above, this work is a first step, and it is quite likely incorporating additional pathways, especially those regulating cell motility, may be necessary to fully decode EMT-metabolism coupling. For instance, the RHO-ROCK signaling network regulates the transition of cancer cells from collective migration of E/M cells to individual migration of amoeboid cells [13,46]. Inclusion of RHO-ROCK signaling could provide a detailed understanding of how metabolism is coupled to different modes of cancer cell migration. Overall, the importance of external signaling in our model is in conceptual agreement with a hypothesis by Sciacovelli and Frezza that, in an adverse TME, metabolic reprogramming drives EMT to allow cells to find favorable metabolic niches [28].

The overall goal of this project is toward understanding all the interrelated aspects of cancer metastasis. Previous studies coupling EMT, stemness, and Notch signaling have shown therapy resistance and increased metastatic potential are associated with stem-like hybrid E/M cells [47–49]. Furthermore, these couplings also resulted in unexpected behaviors such as the co-localization of hybrid E/M cells [47] and a tunable stemness window [48]. Studying individual gene regulatory network modules, even in the presence of signals, is unable to give a thorough understanding of the network properties. Therefore, multiple modules and their crosstalk should be studied concurrently to understand the correlation between cancer traits and potentially identify key regulators.

## Acknowledgements

This work was supported by National Science Foundation by sponsoring the Center for Theoretical Biological Physics – award PHY-2019745 (JNO, HL) and by awards PHY-1605817 (HL), PHY-2210291 (JNO), and PHY-1522550 (JNO, MG). JNO is a CPRIT Scholar in Cancer Research. MG was also supported by the NSF GRFP-1842494.

## Notes

### Competing Interest Statement

The authors have declared no competing interest.

