## Supplementary info for "Decoding the coupled decision-making of the epithelial-mesenchymal transition and metabolic reprogramming in cancer"

### 1 Model

#### 1.1 Regulation modeled by shifted Hill function

For the cases in which transcription factors are regulating other pieces or there is indirect miRNA regulation (e.g., $\mu_{34}$ regulating ROS) then the regulation is modeled as a shifted Hill function

$$H(X, X_0, n, \lambda) = \lambda + \frac{1. - \lambda}{1. + (X/X_0)^n} \tag{1}$$

Where the transcription factor X is regulating Y. $X_0$ is the threshold of X when the regulation becomes stronger. The cooperativity, n, represents the sensitvity to X. The foldchange ($\lambda$) represents the amount of regulation and can be $\lambda < 1$, $\lambda = 1$, or $\lambda > 1$ representing inhibition, no regulation, or activation. Therefore maximum inhibition occurs when $\lambda = 0$ but there is no upper bound on activation.

#### 1.2 miRNA regulation

We utilize the framework for miRNA regulation developed by Lu and collaborators described in detail in Ref.[1]. This framework was developed to model the binding of miRNA and mRNA to form an miRNA-mRNA complex and the subsequent translation of a protein. The miRNA concentration is $\mu$, the treshold of miRNA regulation is $\mu_0$, the mRNA concentration is m, the number of mRNA binding sites for miRNA is n, and the protein concentration is B. The reaction is assumed to occur at steady state and the binding/unbinding of miRNA and mRNA is assumed to be much faster than production/degredation of the protein.

Any number of binding sites may be occupied resulting in $C_i^n$ possible combinations when $i$ binding sites are occupied, where

$$C_i^n = \frac{n!}{i!(n-i)!} \tag{2}$$

When $i$ binding sites are occupied, then

$$[m_i] = mM_n^i(\mu) \tag{3}$$

where

$$M(i, n, \mu, \mu_0) = \frac{\frac{\mu}{\mu_0}^i}{1 + \frac{\mu}{\mu_0}^n} \tag{4}$$

leading to the total translation rate

$$mL(\mu) = \sum_{i=0}^{n} l_i C_i^n M_i^n(\mu, \mu_0) \tag{5}$$

the active mRNA degredation rate

$$mY_m(\mu) = \sum_{i=0}^{n} y_{mi} C_i^n M_i^n(\mu, \mu_0) \tag{6}$$

and the active miRNA degredation rate

$$mY_\mu(\mu) = \sum_{i=0}^{n} i y_{\mu i} C_i^n M_i^n(\mu, \mu_0) \tag{7}$$

Therefore, the dynamics of miRNA-mRNA binding complex and trascribed protein B are modeled by the following equations,

$$\frac{d\mu}{dt} = g_\mu - mY_\mu(\mu) - k_\mu \mu \tag{8}$$

$$\frac{dm}{dt} = g_m - mY_m(\mu) - k_m m \tag{9}$$

$$\frac{dB}{dt} = g_B mL(\mu) - k_B B \tag{10}$$

##### 1.3 Competition in metabolism model

In the metabolic circuit, AMPK and Hif1 competitively regulate the level of mtROS

$$C^{comp}_{Rmt}(\gamma, g_n, h, h^0_{hr_{mt}}, n_{hr_{mt}}, A, A^0_{ar_{mt}}, n_{ar_{mt}} = \frac{\gamma(g_n + (\frac{A}{A^0_{ar_{mt}}})^{n_{ar_{mt}}})}{1 + (\frac{h}{h^0_{hr_{mt}}})^{n_{hr_{mt}}} + (\frac{A}{A^0_{ar_{mt}}})^{n_{ar_{mt}}}} \tag{11}$$

and noxROS

$$C^{comp}_{Rnox}(g_0, h, h^0_{hr_{nox}}, n_{hr_{nox}}, g_1, A, A_0, g_2, n_{ar_{nox}}) = \frac{g_0 + g^h_{r_{nox}}(\frac{h}{h^0_{hr_{nox}}})^{n_{hr_{nox}}} + g^a_{r_{nox}}(\frac{A}{A^0_{ar_{nox}}})^{n_{ar_{nox}}}}{1 + (\frac{h}{h^0_{hr_{nox}}})^{n_{hr_{nox}}} + (\frac{A}{A^0_{ar_{nox}}})^{n_{ar_{nox}}}} \tag{12}$$

##### 1.4 The coupled EMT-metabolism network

$$\frac{dZ}{dt} = g_Z m_z L(\mu_{200}, \mu_{200_0}, n_{\mu_{200}}) - k_Z Z \tag{13}$$

$$\frac{m_Z}{dt} = g_{m_Z} H(Z, Z_{0,m}, n_{Z,m}, \lambda_{Z,m}) H(S, S_{0,m}, n_{s,m}, \lambda_{S,m}) H(A, A_{0,m}, n_{A,m}, \lambda_{A,m}) - m_Z Y_m(\mu, \mu_0, n_\mu) - k_{m_Z} m_Z \tag{14}$$

$$\frac{S}{dt} = g_S m_S L(\mu_{34}, \mu_{34_0}, n_{\mu_{34}}) - k_S S \tag{15}$$

$$\frac{m_S}{dt} = g_{m_S} * H(S, S_{0,ms}, n_{sms}, \lambda_{Sms}) H(I, I_{0m}, n_{Im}, \lambda_{Im}) H(h, h_{0ms}, n_{hms}, \lambda_{hms}) H(A, A_{0ms}, n_{Ams}, \lambda_{Ams}) - m_s Ym(u3, u3_0, n_{u3}) - k_{ms} m_S \tag{16}$$

$$\frac{\mu_{200}}{dt} = gu * H(Z, Z_{0u}, n_{zu}, \lambda_{zu}) H(S, S_{0u}, n_{su}, \lambda_{Su}) H(h, h_{0u}, n_{hu}, \lambda_{hu}) H(A, A_{0u}, n_{Au}, \lambda_{Au}) - m_Z * Yu(u, u_0, n_u) - m_H * Yu(u, u_0, n_{uh}) - k_u * u \tag{17}$$

$$\frac{\mu_{34}}{dt} = g_{u3} * H(Z, Z_{0u3}, n_{zu3}, \lambda_{zu3}) * H(S, S_{0u3}, n_{su3}, \lambda_{Su3}) - m_S * Yu(u3, u3_0, n_{u3}) - k_{u3} * u3 \tag{18}$$

$$\frac{mh}{dt} = g_{mh} * H(A, A_{0ah}, n_{ah}, \lambda_{ah}) - k_{mh} * m_h * H(h, h_{0hh}, n_{hh}, \lambda_{hh}) H(R, R_{0rh}, n_{rh}, \lambda_{rh}) - m_h * Y_m(\mu, \mu_0, n_{\mu,h}, y_{mih}) \quad (19)$$

$$\frac{h}{dt} = g_h m_h L(\mu, \mu_0, n_{\mu,h}, l_{ih}) - k_h h \quad (20)$$

$$\frac{A}{dt} = g_a H(R, R_{0ra}, n_{ra}, \lambda_{ra}) H(h, h_{0ha}, n_{ha}, \lambda_{ha}) H(A, A_{0aa}, n_{aa}, \lambda_{aa}) - k_a * A \quad (21)$$

$$R = R_{mt} + R_{nox} \quad (22)$$

$$\frac{R_{nox}}{dt} = g_{rn} * C^{comp}_{R_{nox}}(g_n, h, h_{0hrn}, n_{hrn}, g_1, A, A_{0rn}, g_2, n_{arn}) - k_{rn} * R_{nox} H(\mu_{34}, n_{u30rn}, n_{3n}, \lambda_{3n}) \quad (23)$$

$$\frac{R_{mt}}{dt} = g_{rm} H(A, A_{0aR}, n_{ar}, \lambda_{ar}) * C^{comp}_{R_{mt}}(\gamma, g_n, h, h_{0hrm}, n_{hrm}, A, A_{0rm}, n_{arm}) - k_{rm} * R_{mt} H(\mu_3, n_{u30rm}, n_{3m}, \lambda_{3m}) \quad (24)$$

##### 1.5 Parameter determination for coupled EMT-metabolic network

The values of parameters for the core EMT are given by Tables 1-1.5, and the parameters for the core metabolic network are given by Table 4. We define the parameters based on an extensive literature search. For example, regarding EMT regulating metabolism, ROS levels are increased by via downregulating the NRF2-dependent antioxidant capability [2, 3, 4] , downregulating SOD2 [5], or upregulating the p53 pathway [6, 7]. Next, family members can either upregulate or downregulate Hif1 expression [9] . While miR-429 upregulates HIF-1, both miR-200b [10] and miR-200c [11] downregulate HIF-1 expression. We focus on the negative feedback loop which seems to be present in a larger portion of the miR-200 family members [9, 12] . Regarding metabolism regulating EMT, HIF-1 inhibits miR-200b through upregulation of the HIF-1 downstream target ASCL2 [10] . Therefore, there is a mutual inhibitory feedback loop between and HIF-1. Additionally, HIF-1 can directly upregulate SNAIL [13], while AMPK represses the production of SNAIL [14] by activating FOXO3. Similarly, AMPK suppresses ZEB2 by activating FOXO1 [15, 16] . Additionally, CREB, after being activated by AMPK via phosphorylation, can transcribe resulting in the upregulation of [17, 18, 19, 20, 21].

Any parameter in Table 1.5 with a citation is an experimentally derived value that we are using. Any parameter without a reference is estimated based on what is known of the system and other parameters. The values for the L, Ym, and Yu functions are estimated and set in ranges to ensure the behavior mimics biological systems.

Since Hif1 is regulated by miRNA $\mu_{200}$, it was required to also include Hif1 messenger RNA which was not inlcuded in the original circuit studied by Yu et al [22]. This ensured new parameters for Hif1, and Hif1 mRNA, were required therefore we went through a range of parameters and found the parameters that gave the most similar result with or without $\mu_{200}$ regulating Hif1. See Tables 4 and 1.5 for parameter values.

The parameters for the coupled links are defined within a range from inactive to fully active. Unless specified, all crosstalks are assumed to be in the inactive state. For crosstalks modeled as shifted Hill functions, the value of the foldchange is set to one ($\lambda_{crosstalk} = 1$). If the crosstalk for between $\mu_{200}$ and Hif1 is inactive ($\mu_{200} - |$ HIF-1), the production of HIF-1 is decoupled from $\mu_{200}$. This decoupling reflects how HIF-1 production was initially defined in the metabolic network of Yu and collaborators [22]. To mathematically model this decoupling, the value of $L_H(\mu_{200}) = 1$, $Y_{m,H}(\mu_{200})$=0, and $Y_{\mu,H}(\mu_{200}) = 0$.

| | molecules/hr | | $\text{hr}^{-1}$ | | | | | | molecules |
|---|---|---|---|---|---|---|---|---|---|
| $g_z$ | 100 | $k_z$ | 0.1 | $\lambda_{zu}$ | 0.1 | $\text{n}_{\mu_{200}}$ | 6 | S0Z | 180000 |
| $g_{mz}$ | 11 | $k_{m_z}$ | 0.5 | $\lambda_{Su}$ | 0.1 | $n_{u3}$ | 2 | I | 50000 |
| $g_S$ | 100 | $k_S$ | 0.125 | $\lambda_{Im}$ | 10 | $n_{zu}$ | 3 | $u_{200_0}$ | 10000 |
| $g_{ms}$ | 90 | $k_{ms}$ | 0.5 | $\lambda_{Sms}$ | 0.1 | $n_{su}$ | 2 | u30 | 10000 |
| $gu$ | 2100 | $k_u$ | 0.05 | $\lambda_{Zu3}$ | 0.2 | $n_{su3}$ | 1 | $Z_{0u}$ | 220000 |
| $g_{u3}$ | 1350 | $k_{u3}$ | 0.05 | $\lambda_{Su3}$ | 0.1 | $n_{s,m}$ | 2 | $S_{0,ms}$ | 200000 |
| | | | | $\lambda_{Z,m}$ | 7.5 | $n_{Z,m}$ | 2 | $S_{0u}$ | 180000 |
| | | | | $\lambda_{S,m}$ | 10. | $n_{Im}$ | 2 | $I_{0m}$ | 50000 |
| | | | | | | $n_{sms}$ | 1 | $Z_{0,m}$ | 25000 |
| | | | | | | $n_{zu3}$ | 2 | $Z_{0u3}$ | 600000 |
| | | | | | | | | $S_{0u3}$ | 300000 |

Table 1: Parameters of the core EMT network - u200/ZEB/u34/SNAIL

| n( # of miRNA binding sites) | i=0 | i=1 | i=2 | i=3 | i=4 | i=5 | i=6 |
|---|---|---|---|---|---|---|---|
| $l_i$ | 1 | 0.6 | 0.3 | 0.1 | 0.05 | 0.05 | 0.05 |
| $y_{mi}$ | 0 | 0.04 | 0.2 | 1.0 | 1.0 | 1.0 | 1.0 |
| $y_{\mu i}$ | 0 | 0.005 | 0.05 | 0.5 | 0.5 | 0.5 | 0.5 |

Table 2: Parameters for regulation by $\mu_{200}$ in the EMT network

| n( # of miRNA binding sites) | i=0 | i=1 | i=2 |
|---|---|---|---|
| $l_i$ | 1 | 0.6 | 0.3 |
| $y_{mi}$ | 0 | 0.04 | 0.2 |
| $y_{\mu i}$ | 0 | 0.005 | 0.05 |

Table 3: Parameters for regulation by $\mu_{34}$ in the EMT network

| | nM/hr | | $\mu M$/min | | $\text{hr}^{-1}$ | | $\text{min}^{-1}$ | | nM | | $\mu M$ | | |
|---|---|---|---|---|---|---|---|---|---|---|---|---|---|
| $g_a$ | 30 | $g_{rn}$ | 40 | $k_a$ | 0.2 | $k_{rn}$ | 5 | $H_{0hh}$ | 80 | $R_{0rh}$ | 300 | $\lambda_{rh}$ | 0.2 |
| $g_h$ | 1.5 | $g_{rm}$ | 150 | $k_h$ | 1.75 | $k_{rm}$ | 5 | $A_{0ah}$ | 250 | $R_{0ra}$ | 100 | $\lambda_{hh}$ | 0.1 |
| $g_{mh}$ | 10 | | | $k_{mh}$ | 0.143 | | | $A_{0aR}$ | 350 | | | $\lambda_{ah}$ | 0.1 |
| | | | | | | | | $H_{0ha}$ | 250 | | | $\lambda_{ar}$ | 0.25 |
| | | | | | | | | $A_{0aa}$ | 350 | | | $\lambda_{ra}$ | 8. |
| | | | | | | | | $h_{0hrm}$ | 200 | | | $\lambda_{ha}$ | 0.1 |
| | | | | | | | | $h_{0hrn}$ | 250 | | | $\lambda_{aa}$ | 0.2 |
| | | | | | | | | $A_{0rn}$ | 150 | | | $n_{rh}$ | 4 |
| | | | | | | | | $A_{0rm}$ | 150 | | | $n_{hh}$ | 4 |
| | | | | | | | | | | | | $n_{ah}$ | 1 |
| | | | | | | | | | | | | $n_{ar}$ | 2 |
| | | | | | | | | | | | | $n_{ra}$ | 4 |
| | | | | | | | | | | | | $n_{ha}$ | 1 |
| | | | | | | | | | | | | $n_{aa}$ | 2 |
| | | | | | | | | | | | | $g_n$ | 0.2 |
| | | | | | | | | | | | | $g_1$ | 5 |
| | | | | | | | | | | | | $g_2$ | 0.2 |
| | | | | | | | | | | | | $\gamma$ | 8 |
| | | | | | | | | | | | | $n_{arm}$ | 4 |
| | | | | | | | | | | | | $n_{hrm}$ | 2 |
| | | | | | | | | | | | | $n_{hrn}$ | 2 |
| | | | | | | | | | | | | $n_{arn}$ | 2 |

Table 4: Parameters of the core metabolic network - AMPK/HIF-1/ROS

| Reg | foldchange | | cooperativity | | Threshold | | References |
|---|---|---|---|---|---|---|---|
| | parameter | value | parameter | value | parameter | value | |
| AS | $\lambda_{AS}$[14] | 0. | nAS | 2 | A0S | 300 | [14] |
| AZ | $\lambda_{AS}$ | | nAZ[15] | 2 | A0Z | 300 | [15, 16] |
| Au | $\lambda_{Au}$[18] | 0.6 | nAu[20] | 1 | A0u | 300 | [17, 18, 19, 20, 21] |
| HS | $\lambda_{HS}$ [13] | 7 | nHS[13] | 2 | H0S | 200 | [13] |
| Hu | $\lambda_{Hu}$[10] | 1.5 | nHu | 1 | H0u | 200 | [9, 12, 10] |
| u3m | $\lambda_{u3m}$[5] | 2 | n3m | 3 | u30m | 10000 | [2, 3, 4, 5, 6, 7] |
| u3n | $\lambda_{AS}$ | | n3n | 2 | u30n | 10000 | [2, 3, 4, 5, 6, 7] |
| uh | | | nuh | 2 | | | [9, 10, 11, 9, 12] |

Table 5: Crosstalk between the EMT and metabolism networks

| $l_i$ | | | $y_{m,i}$ | | | $y_{\mu,i}$ | | |
|---|---|---|---|---|---|---|---|---|
| i=0 | i=1 | i=2 | i=0 | i=1 | i=2 | i=0 | i=1 | i=2 |
| 1 | 0 | 0 | 0 | 0.002 | 0.01 | 0 | 0.001 | 0.009 |
| 1 | 0 | 0 | 0 | 0.01 | 0.5 | 0 | 0.001 | 0.009 |
| 1 | 0 | 0 | 0 | 0.01 | 0.5 | 0 | 0.01 | 0.09 |
| 1 | 0 | 0 | 0 | 2 | 4 | 0 | 0.001 | 0.009 |
| 1 | 0 | 0 | 0 | 2 | 4 | 0 | 0.01 | 0.09 |
| 1 | 0 | 0 | 0 | 2 | 4 | 0 | 0.1 | 0.2 |
| 1 | 0 | 0 | 0 | 2 | 4 | 0 | 0.5 | 1. |
| 1 | 0.2 | 0.2 | 0 | 0.002 | 0.01 | 0 | 0.001 | 0.009 |
| 1 | 0.2 | 0.2 | 0 | 0.01 | 0.5 | 0 | 0.001 | 0.009 |
| 1 | 0.2 | 0.2 | 0 | 0.01 | 0.5 | 0 | 0.01 | 0.09 |
| 1 | 0.2 | 0.2 | 0 | 2 | 4 | 0 | 0.001 | 0.009 |
| 1 | 0.2 | 0.2 | 0 | 2 | 4 | 0 | 0.01 | 0.09 |
| 1 | 0.2 | 0.2 | 0 | 2 | 4 | 0 | 0.1 | 0.2 |
| 1 | 0.2 | 0.2 | 0 | 2 | 4 | 0 | 0.5 | 1. |
| 1 | 0.5 | 0.3 | 0 | 0.002 | 0.01 | 0 | 0.001 | 0.009 |
| 1 | 0.5 | 0.3 | 0 | 0.01 | 0.5 | 0 | 0.001 | 0.009 |
| 1 | 0.5 | 0.3 | 0 | 0.01 | 0.5 | 0 | 0.01 | 0.09 |
| 1 | 0.5 | 0.3 | 0 | 2 | 4 | 0 | 0.001 | 0.009 |
| 1 | 0.5 | 0.3 | 0 | 2 | 4 | 0 | 0.01 | 0.09 |
| 1 | 0.5 | 0.3 | 0 | 2 | 4 | 0 | 0.1 | 0.2 |
| 1 | 0.5 | 0.3 | 0 | 2 | 4 | 0 | 0.5 | 1. |
| 1 | 0.9 | 0.8 | 0 | 0.002 | 0.01 | 0 | 0.001 | 0.009 |
| 1 | 0.9 | 0.8 | 0 | 0.01 | 0.5 | 0 | 0.001 | 0.009 |
| 1 | 0.9 | 0.8 | 0 | 0.01 | 0.5 | 0 | 0.01 | 0.09 |
| 1 | 0.9 | 0.8 | 0 | 2 | 4 | 0 | 0.001 | 0.009 |
| 1 | 0.9 | 0.8 | 0 | 2 | 4 | 0 | 0.01 | 0.09 |
| 1 | 0.9 | 0.8 | 0 | 2 | 4 | 0 | 0.1 | 0.2 |
| 1 | 0.9 | 0.8 | 0 | 2 | 4 | 0 | 0.5 | 1. |

Table 6: Parameters for regulation of HIF-1 by $\mu_{200}$

#### 1.6 Coupled EMT-metabolism network without hybrid phenotypes

to generate a model that is missing the hybrid phenotype we adjusted the following parameters.

For the results labled noEM:
$\lambda_{Su} = 0.85$ and $\lambda_{SZ} = 17$

For the results labeled noWO:
kmh = 0.158, kh = 2.2, and $\gamma = 6$

For the results label noHH: We removed the initial ability to access the WO or EM states.
$\lambda_{Su} = 0.85$, $\lambda_{SZ} = 17$, kmh = 0.158, kh = 2.2, and $\gamma = 6$

The states were confirmed by calculating the nullclines of the EMT and metabolic circuit with the new parameters.

#### 1.7 Coupled EMT-metabolism network with Phenotype stability factors

When introducing the PSFs OVOL and GRHL2 we add four equations to our model that represent the protein and mRNA levels of OVOL and GRHL2. We also must adjust the equations for $\mu_{200}$ to include inhibition by OVOl, Zeb mRNA to include inhibition by OVOL and GRHL2, and ROS to include upregulation by GRHL2. The parameters that have changed or been added are listed in Table 7

$$\frac{m_Z}{dt} = g_{m_Z} H(Z, Z_{0,m}, n_{Z,m}, \lambda_{Z,m}) H(S, S_{0,m}, n_{s,m}, \lambda_{S,m}) H(A, A_{0,m}, n_{A,m}, \lambda_{A,m}) \\ H(O, O_{0,m}, n_{O,m}, \lambda_{O,m}) H(G, G_{0,m}, n_{G,m}, \lambda_{G,m}) - m_Z Y_m(\mu, \mu_0, n_\mu) - k_{m_Z} m_Z \quad (25)$$

$$\frac{\mu_{200}}{dt} = gu * H(Z, Z_{0u}, n_{zu}, \lambda_{zu}) H(S, S_{0u}, n_{su}, \lambda_{Su}) H(h, h_{0u}, n_{hu}, \lambda_{hu}) \\ H(A, A_{0u}, n_{Au}, \lambda_{Au}) H(O, O_{0,u}, n_{O,u}, \lambda_{O,u}) - m_Z * Yu(u, u_0, n_u) - m_H * Yu(u, u_0, n_{uh}) - k_u * u \quad (26)$$

$$\frac{G}{dt} = g_g m_g - k_{gg} G \quad (27)$$

$$\frac{m_g}{dt} = g_{m_g} H(Z, Z_{0,mg}, n_{z,mg}, \lambda_{z,mg}) - k_{m_g} m_g \quad (28)$$

$$\frac{O}{dt} = g_o m_o - k_o O \quad (29)$$

$$\frac{m_o}{dt} = g_{m_o} H(O, o_{0,mo}, n_{o,mo}, \lambda_{o,mo}) H(Z, Z_{0,mo}, n_{z,mo}, \lambda_{z,mo}) H(G, G_{0,mo}, n_{g,mo}, \lambda_{g,mo}) - k_{m_o} m_o \quad (30)$$

When the crosstalks are inactive, we find the PSF stabilized network is mainly in the E/M-W/O state with less than 10% occupancy of the E/M-O state.

| | | | | | | | molecules | | molecules/hr | | $hr^{-1}$ |
|---|---|---|---|---|---|---|---|---|---|---|---|
| $\lambda_{I,S}$ | 16 | $n_{O,\mu}$ | 1 | $O_{0,\mu}$ | 250000 | $g_g$ | 200 | $k_O$ | 0.1 |
| $\lambda_{O,\mu}$ | 0.1 | $n_{O,Z}$ | 1 | $O_{0,Z}$ | 25000 | $g_{mg}$ | 22 | $k_{mO}$ | 0.5 |
| $\lambda_{O,Z}$ | 0.1 | $n_{O,O}$ | 2 | $O_{0,O}$ | 25000 | $g_O$ | 200 | $k_G$ | 0.1 |
| $\lambda_{O,O}$ | 0.1 | $n_{Z,O}$ | 1 | $Z_{0,O}$ | 10000 | $g_{mO}$ | 22 | $k_{mG}$ | 0.5 |
| $\lambda_{Z,O}$ | 0.5 | $n_{Z,G}$ | 3 | $Z_{0,G}$ | 10000 | | | | |
| $\lambda_{Z,G}$ | 0.5 | $n_{G,Z}$ | 1 | $G_{0,Z}$ | 25000 | | | | |
| $\lambda_{G,Z}$ | 0.1 | $n_{G,O}$ | 2 | $G_{0,O}$ | 25000 | | | | |
| $\lambda_{G,O}$ | 0.7 | $n_{G,R_{mt}}$ | 1 | $G_{0,R_{mt}}$ | 25000 | | | | |
| $\lambda_{G,R_{mt}}$ | 0.26 | $n_{G,R_{nox}}$ | 1 | $G_{0,R_{nox}}$ | 25000 | | | | |
| $\lambda_{G,R_{nox}}$ | 0.25 | | | | | | | | |

Table 7: Coupled EMT-metabolic circuit with PSF function parameters

### 2 Methods

#### 2.1 Solving the model

Starting from a set of 1000 distinct random conditions sampled from a uniform distrubtion (ranges for each component in Table 8), we solve the model with the Euler method. For each initial condition we use a timestep of dt=0.1 with relaxation time of 1000 hr, and the values are assumed to have converged at the end of the simulation. The results presented here for each foldchange value, except for $\mu_{200} - |Hif1$, are the average of the results generated from these 1000 initial conditions. The regulation of Hif1 by $\mu_{200}$ has nine distinct parameter values that can be modified, so the results for each initial condition are individually analyzed and then combined. If the parameters for this crosstalk are changed the quantitative results may be misleading, therefore we focus on qualitative results for this crosstalk.

| parameter | range |
|---|---|
| z | [0,700000] |
| mz | [0,2000] |
| S | [0,250000] |
| ms | [0,1000] |
| $\mu_{200}$ | [0,25000] |
| $\mu_{34}$ | [0,20000] |
| A | [0,1000] |
| h | [0,1000] |
| mh | [0,1000] |
| Rmt | [0,1000] |
| Rnox | [0,1000] |

Table 8: Ranges for initial conditions

#### 2.2 Nullclines

To confirm the stability of these states we calculated the continuity and nullclines of the systems using PyDSTool[23]. We calculated the nullclines for the system with inactive crosstalks allowing us to independently model the EMT and metabolic networks. The maximum search was set to $1e + 4$, the error was set to $1e - 10$, maximum step size was $1e + 2$, minimum step size was $1e = 0$, and all bifurcation points were found.

#### 2.3 Coupled state classification

To determine the states we compare to the coupled circuits with inactive crosstalks and original snail input (I=50000). The coupled network excluding the hybrid states were also compared to the inactive crosstalks.

Because the coupled network with the PSFs generates the hybrid state with a different expression profile than the inactive circuit, we generated a new set of gene profiles corresponding to the E/M, E, and M states.

The state is then calculated by determining which of the 9 possible coupled states of the inactive system is closest to the coupled result

$$(d_i^C)^2 = min(\{\sum_j (log_{10}(\frac{x_{i,j}^C}{x_{k,j}^{IA}}))^2 : j \in \{H, A, Z, ...\}, k \in \{E\text{-}W, E\text{-}O, E\text{-}W/O, E/M\text{-}O...\}\}) \quad (31)$$

Whichever state (k) corresponds to the minimum distance for the *ith* generated expression profile ($d_i^C$) is then the assumed state. By taking the square of the log before summing we ensure that deviations from the gene expression profile in opposite directions do not cancel out.

#### 2.4 Phase planes and calculating up/downregulations

To generate the how the possible coupled states change as the regulation changes we determine this by looking at a phase plane of the results. For all regulatory crosstalks that are modeled by a shifted Hill function (all crosstalks except miRNA regulation of Hif-1 by $\mu_{200}$), all 1000 initial conditions at a specific value of the foldchange for that crosstalk $\lambda_{A->|B}$ are classified as one of the nine possible coupled states. For $\mu_{200} - |$ Hif1, the silencing value takes the place of the foldchange value. Thus, we have transformed potentially 1000 different results to between 1 and 9 different results (coupled states). So for each regulatory value ($\lambda_{A->|B}$ or $P_H(\mu_{200})$ we have a set of possible coupled states. Each set is identified by a unique color (with each set of possible coupled states noted either directly on the plot or in a nearby legend). As our focus is on the presence and stability of the E/M-W/O state, we can overlay all plots with black or red dots. The black dots overlaying a plot reference the regulatory parameters for which the E/M-W/O coupled state is possible, while the red dots overlaying a plot are in the regions of regulatory parameters where only the E/M-W/O coupled state exists (i.e., all other coupled states are suppressed).

To determine if a state is up/downregulated relative to the inactive circuit we simply calculate the fraction of initial conditions leading to the E/M-W/O state for the coupled circuit compared to the inactive system. For the coupled network and the coupled network excluding the hybrid states, the inactive network is the tristable network with all crosstalks inactive. For the coupled network including the PSFs, the inactive network is only able to access the E/M-W/O and E/M-O states.

#### 2.5 Silencing Hif-1 mRNA

As the miRNA regulation has three sets of possible parameters changing (l,$y_m$, and $y_\mu$), we utilize the silencing function to incorporate all changing parameters into a single variable.

The silencing function is defined as

$$P_H(\mu_{200}) = 1 - \frac{L(\mu_{200})}{1 + Y_m(\mu_{200})/k_{mh}} \quad (32)$$

We use the degradation rate of the Hif-1 mRNA($k_{mh} = 0.143$).

We calculate the silencing value of each initial condition for a distinct set of parameter.

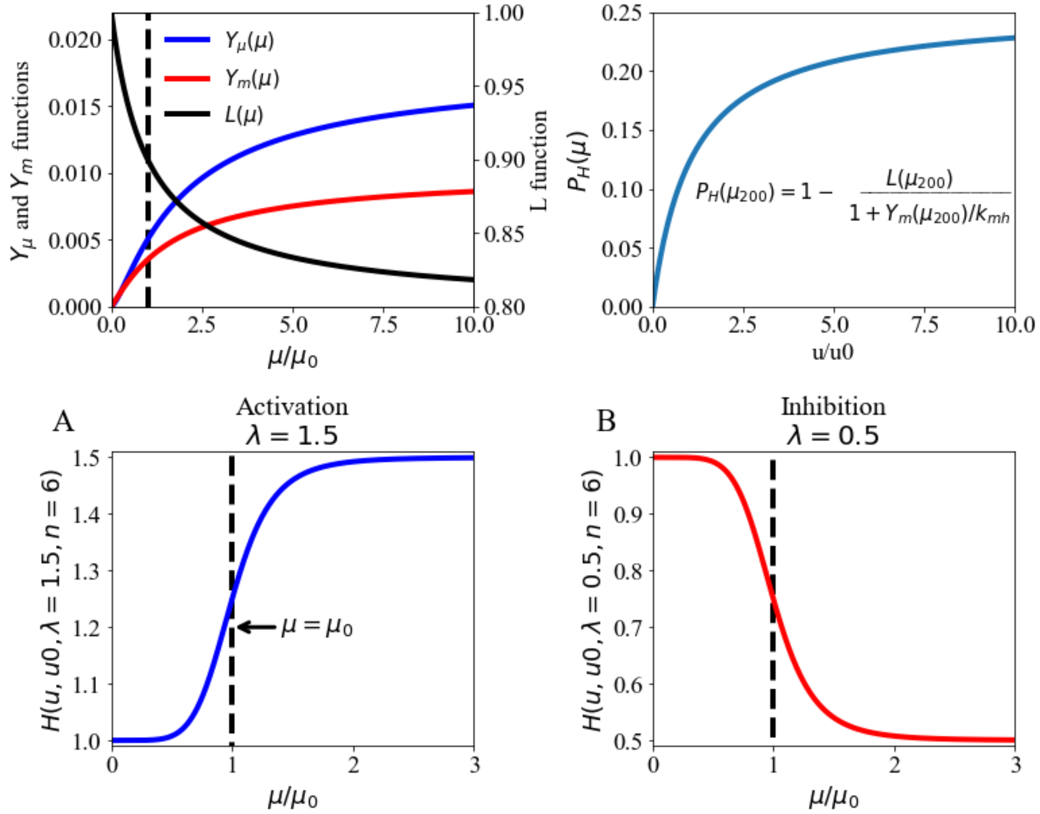


Figure 1: (A) The miRNA regulatory functions for $\mu_{200}$ inhibiting Hif-1 (n=2, $\mu_0 = 10000$, $y_\mu$=[0,0.001,0.009], $y_m$=[0,0.002,0.01], and $l$=[1,0.9,0.8] ). Left axis shows the values for $Y_\mu$ and $Y_m$, degradation rate of the miRNA and mRNA, respectively. Right axis shows values for $L$, translation rate. (B) The silencing function for $\mu_{200} - |Hif1$ ($k_{mh} = 0.143$, n=2, $\mu_0 = 10000$, $y_m$=[0,0.002,0.01], and $l$=[1,0.9,0.8]). (C) The shifted hill functions model transcriptional regulation for n=6 and $\mu_0 = 10000$, and transcriptional activation with $\lambda = 1.5$. (D) Same as (C) but for transcriptional inhibition with $\lambda = 0.5$.

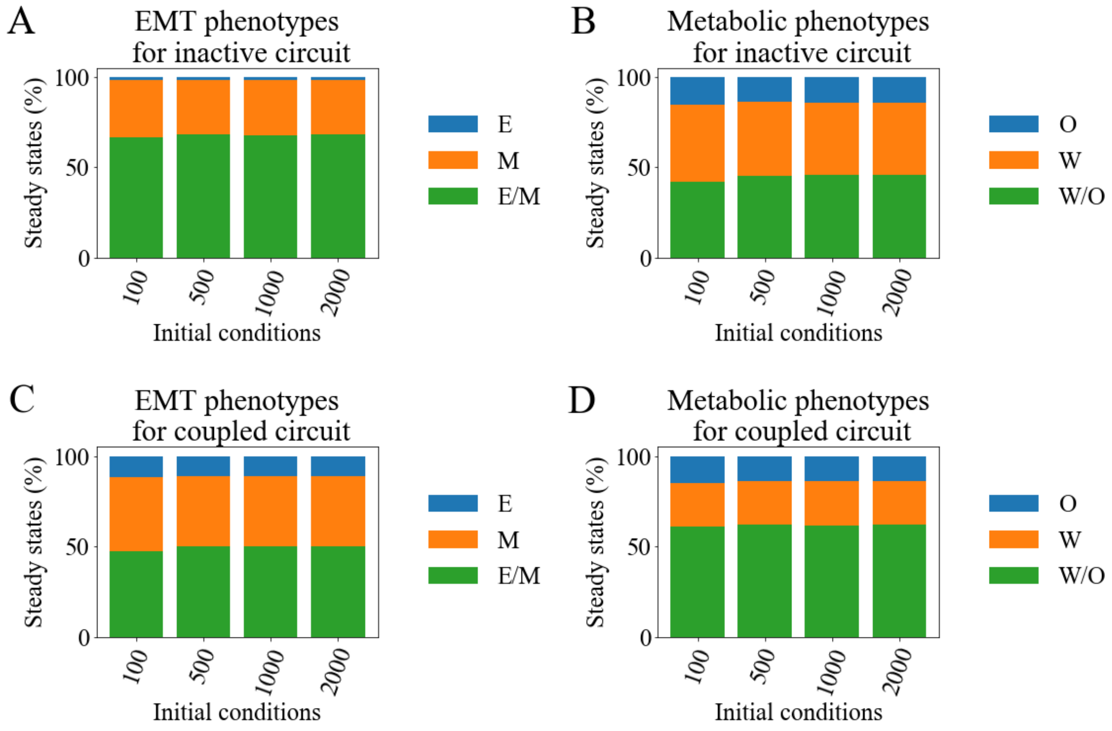


Figure 2: (A) The percent of initial conditions of the coupled, inactive network leading to E (blue), M (orange), or E/M (green) steady states for 100, 500, 1000, and 2000 random initial conditions. (B) Same as A but for the metabolic network. (C) Same as A but for a coupled network ($\lambda_{AMPK-|Snail}$=0.95,$\lambda_{AMPK-|Zeb} =$ 0.95 , $\lambda_{Hif1-|\mu_{200}} = 0.9$, $\lambda_{\mu_{34}->mtROS} = 0.7$, $\lambda_{\mu_{34}->noxROS} = 0.7$, $\lambda_{Hif1->Snail} = 1.1$, and $\lambda_{Ampk->\mu_{200}} =$ 1.01). (D) Same as (C) but for metabolic network.

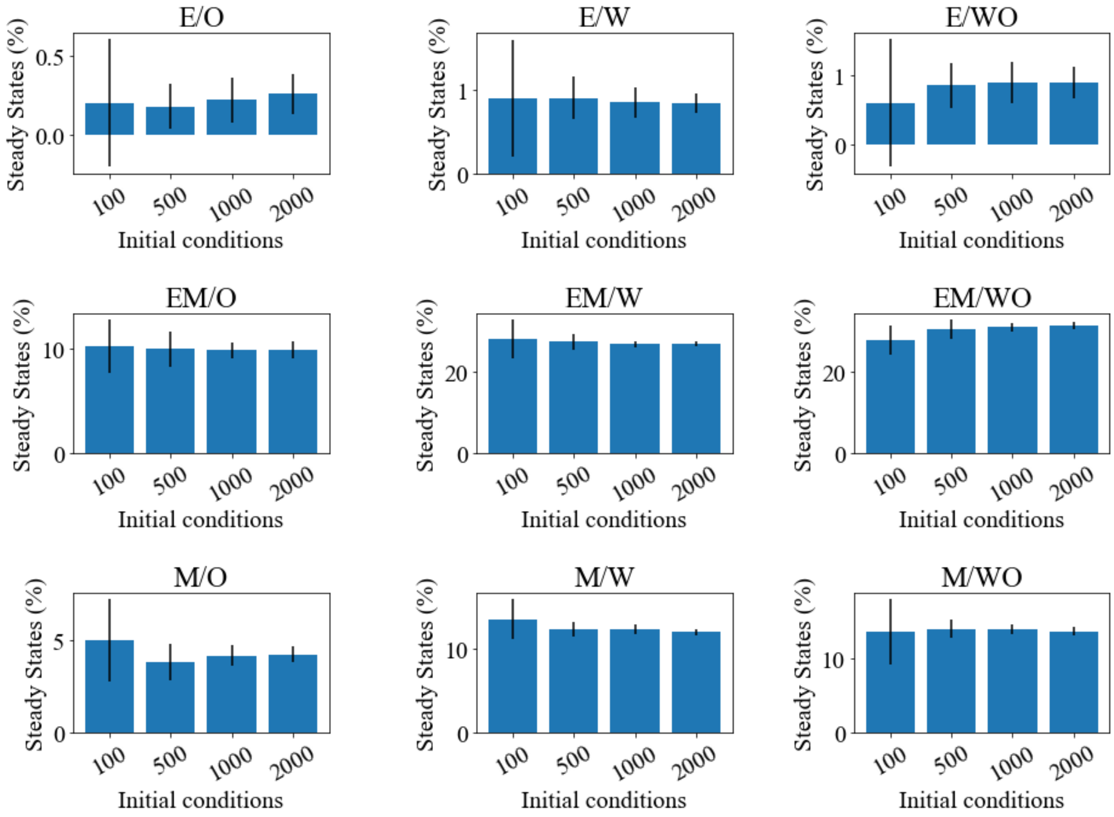


Figure 3: The percent of initial conditions leading to the coupled steady states for 100, 500, 1000 and 2000 sets of random initial conditions for the coupled, inactive network. Corresponds with Fig. 2A-B

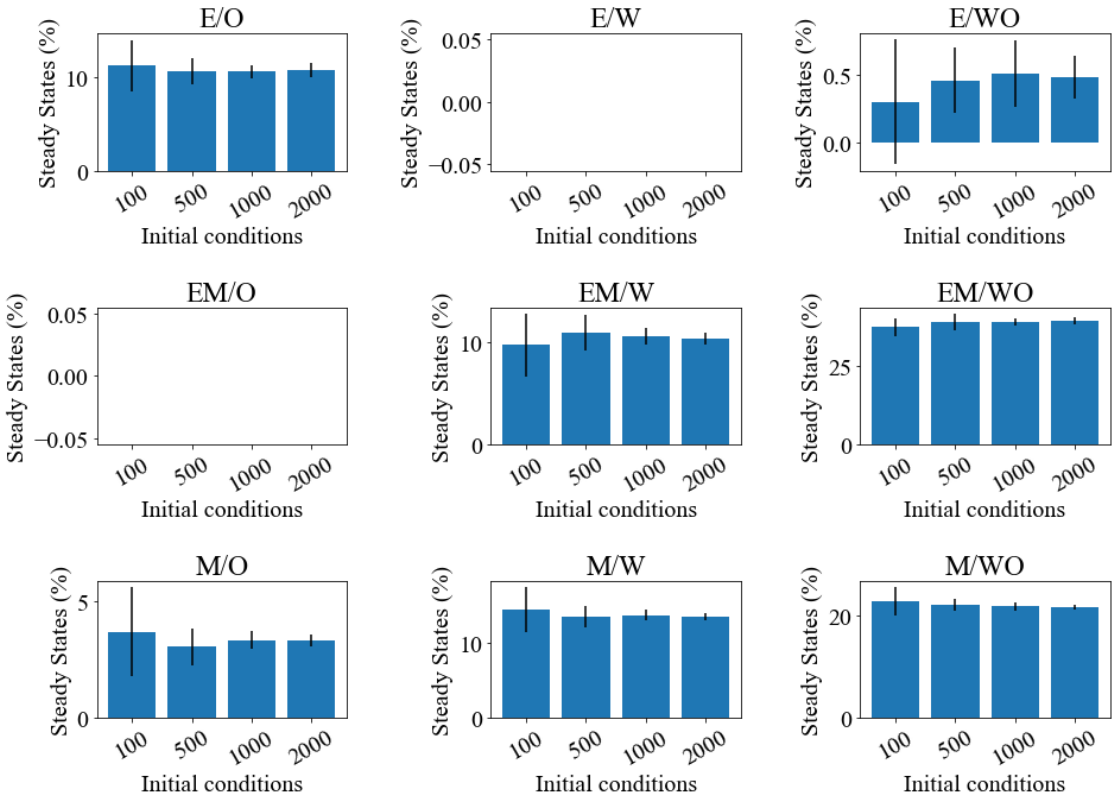


Figure 4: The percent of initial conditions leading to the coupled steady states for 100, 500, 1000 and 2000 sets of random initial conditions for the coupled network ($\lambda_{AMPK-|Snail}$=0.95,$\lambda_{AMPK-|Zeb} = 0.95$ , $\lambda_{Hif1-|\mu_{200}} = 0.9$, $\lambda_{\mu_{34}->mtROS} = 0.7$, $\lambda_{\mu_{34}->noxROS} = 0.7$, $\lambda_{Hif1->Snail} = 1.1$, and $\lambda_{Ampk->\mu_{200}} = 1.01$). Corresponds with Fig. 2C-D

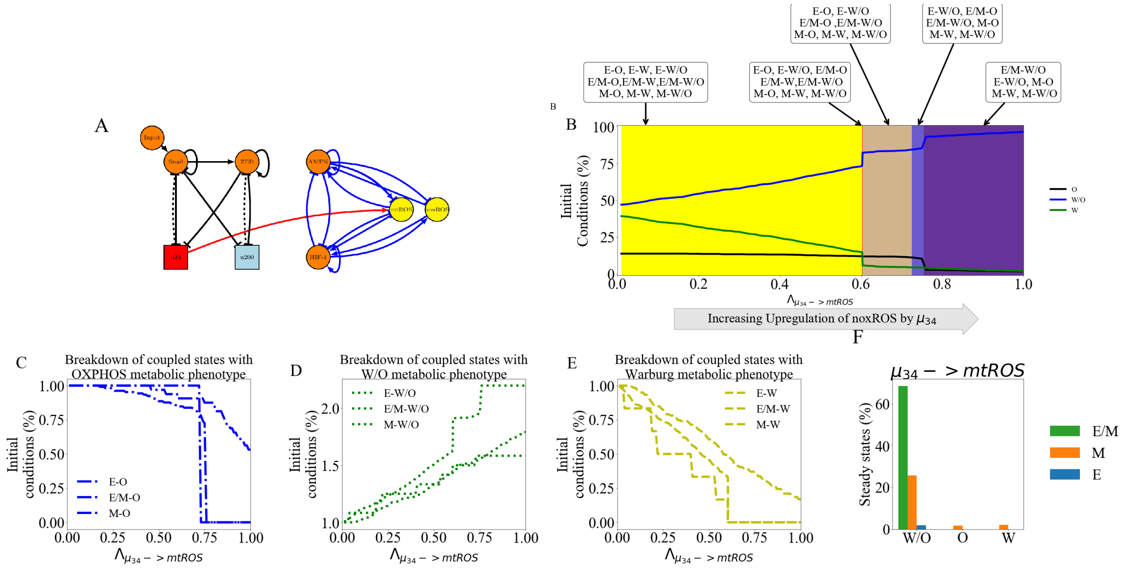

Figure 5: $\mu_{34}- >$mtROS is only active crosstalk. (A) The network showing the active crosstalk. (B) The possible sets of coupled steady states as the regulation increases. (C) The fraction of initial conditions at a particular level of regulation ($\lambda$) compared to the inactive state ($\lambda = 1$) that lead to the E-O, E/M-O, or M-O coupled states. (D) The same as (C) except for the coupled states with metabolic phenotype W/O. (E) The same as (C) except for the coupled states with metabolic phenotype W. (F) The percent of initial conditions leading to W/O, O, and W phenotypes broken down by coupled EMT phenotype. As mtROS is upregulated by $\mu_{34}$, the W and O states are slowly suppresed until the system is nearly 100 W/O phenotype.

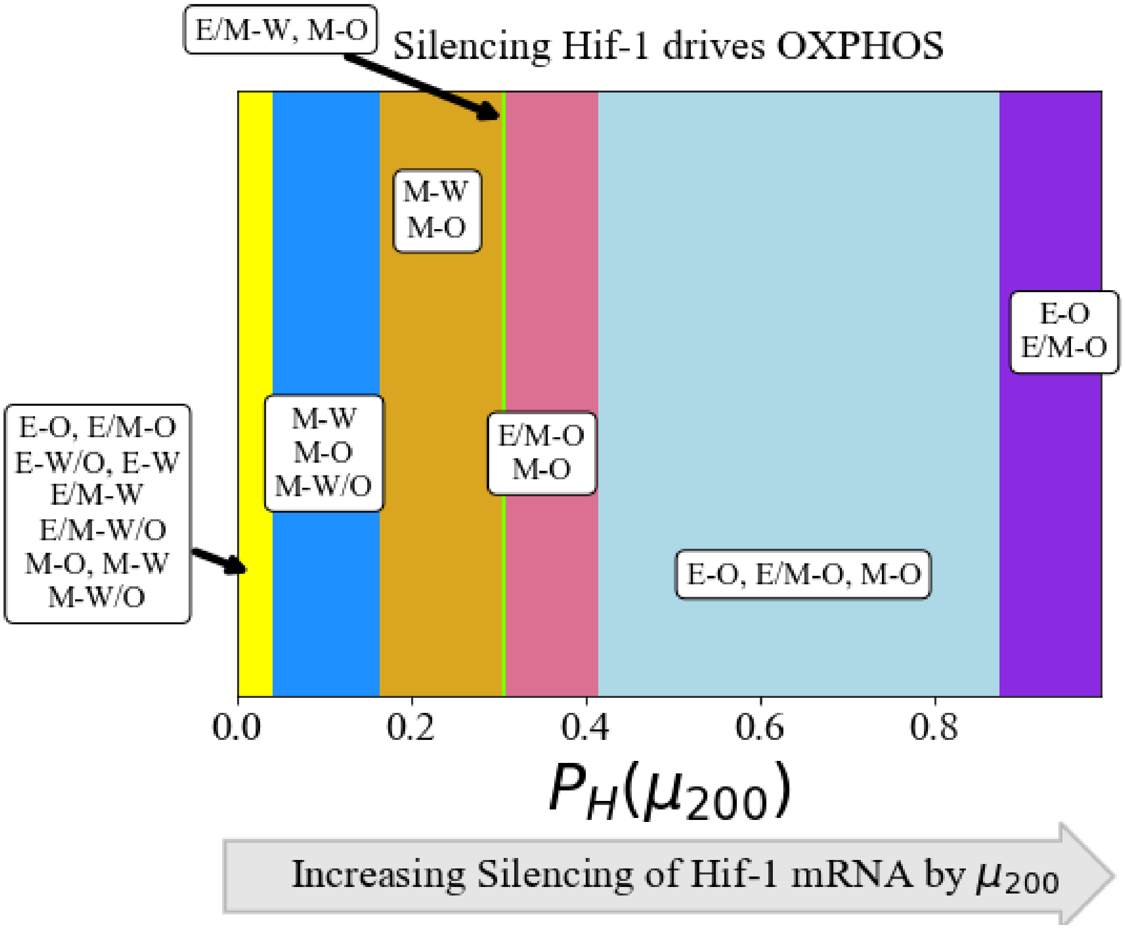


Figure 6: The coupled phenotypes associated with increased silencing of the HIF-1 mRNA by $\mu_{200}$ . Both the EMT and metabolic networks are affected by the $\mu_{200}$ mediated regulation of HIF-1, and the E/M-W/O state is suppressed if any fraction of silencing occurs. At minimal silencing ($P_H(\mu_{200})$( near 1) only the coupled states with M are accessible (M-W, M-O, and M-W/O). Then as $\mu_{200}$-mediated inhibition of HIF-1 increases, the W/O state is suppressed, then the M-W state undergoes partial EMT and becomes the E/M-W state, and after that only the O-associated states are accessible. At complete silencing of the HIF-1 mRNA only the E-O and E/M-O states are accessible.

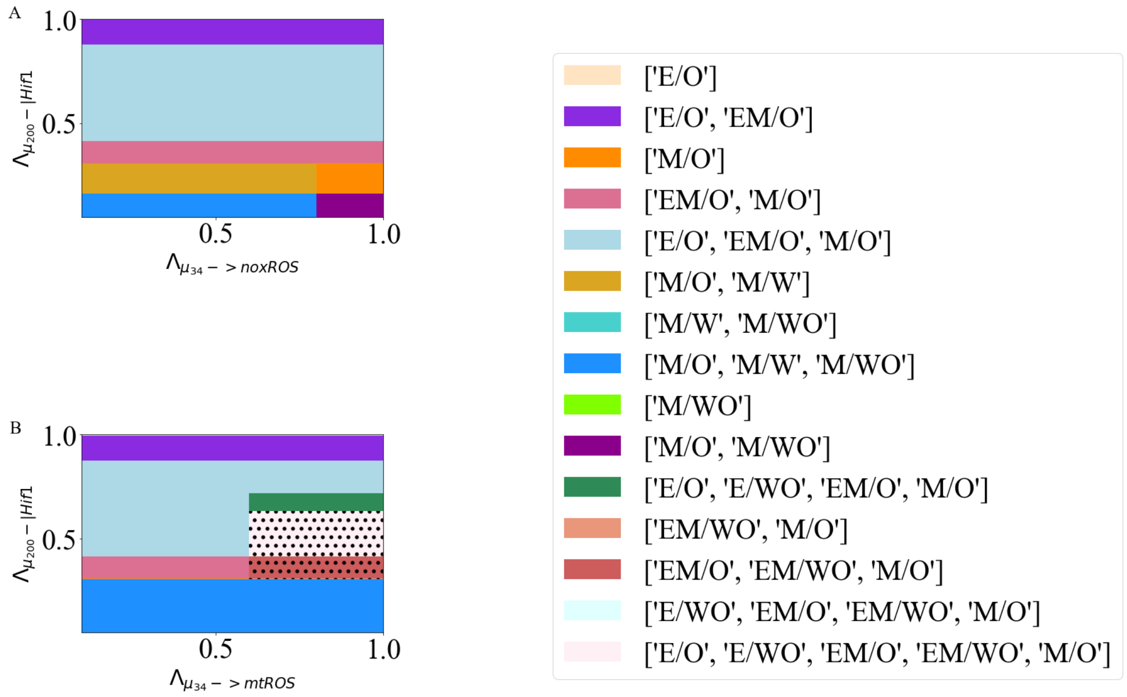


Figure 7: (A) The potential sets of steady states available as the level of noxROS is upregulated (x-axis) and Hif1 is silenced (y-axis). (B) Same as (A) for upregulation of mtROS (x-axis) and Hif1 silencing (y-axis). The legend is for both A and B. The E/M-W/O state is only accessible in the black dotted region of B.

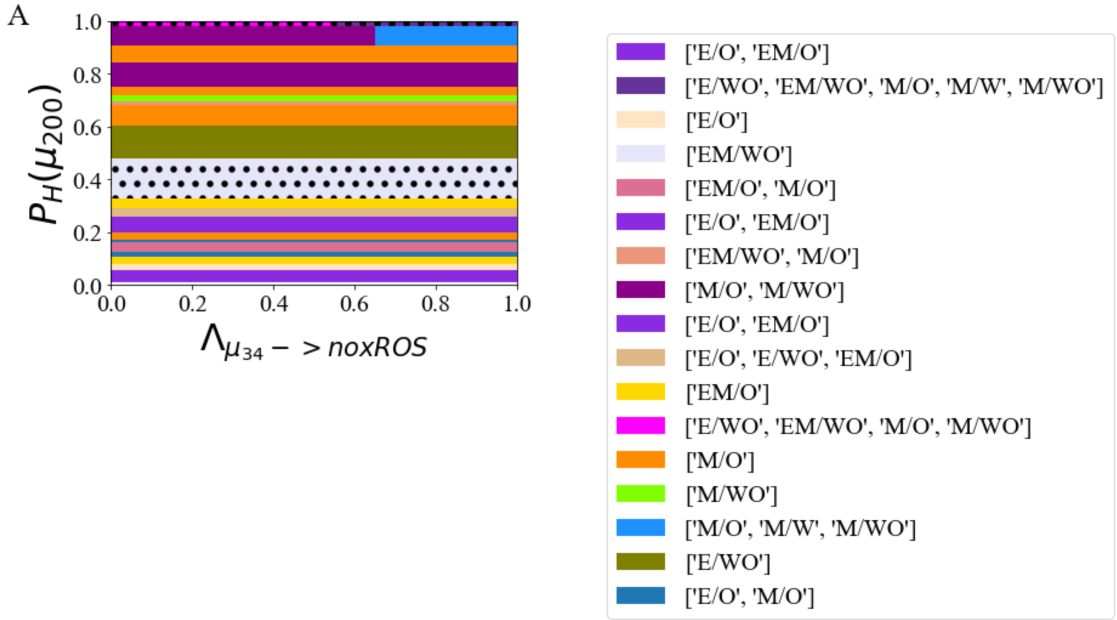


Figure 8: The coupled phases as a function of noxROS upregulation and HIF-1 silencing, when $\mu_{34}->mtROS$, $\mu_{34}->$noxROS, and $\mu_{200}-|$HIF-1. The E/M-W/O state is present for intermediate silencing of HIF-1. The results are minimally reliant on noxROS regulation.

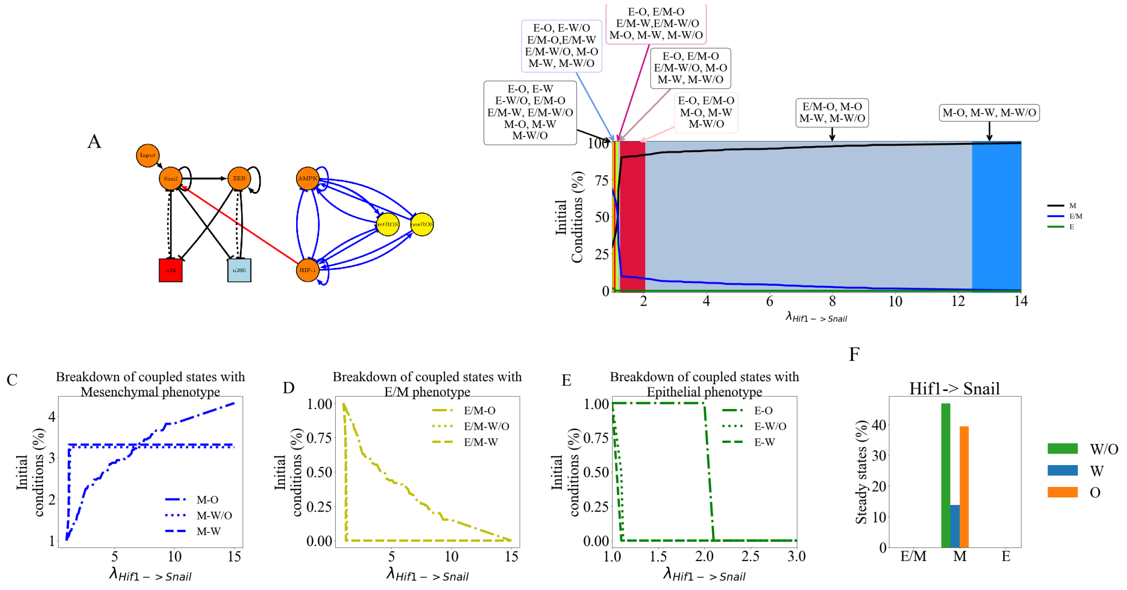


Figure 9: Hif1$->$ Snail is only active crosstalk. (A) The network showing the active crosstalk. (B) The possible sets of coupled steady states as the regulation increases. (C) The fraction of initial conditions at a particular level of regulation ($\lambda$) compared to the inactive state ($\lambda = 1$) that lead to the M-O, M-W/O, or M-W coupled states. (D) The same as (C) except for the coupled states with E/M phenotype. (E) The same as (C) except for the coupled states with E phenotype. (F) The percent of initial conditions leading to E/M, M, and E phenotypes broken down by coupled metabolic phenotype. As Snail is upregulated by Hif-1, the E/M and E states are slowly suppresed until the system is saturated at mesenchymal.

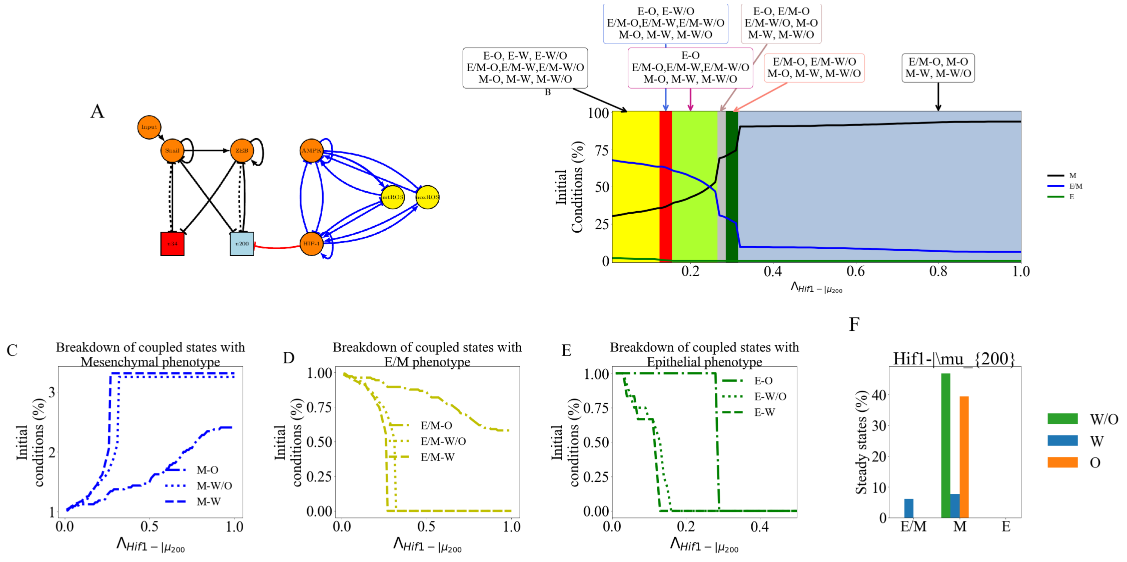


Figure 10: Hif1$-|\mu_{200}$ is only active crosstalk. (A) The network showing the active crosstalk. (B) The possible sets of coupled steady states as the regulation increases. (C) The fraction of initial conditions at a particular level of regulation ($\lambda$) compared to the inactive state ($\lambda = 1$) that lead to the M-O, M-W/O, or M-W coupled states. (D) The same as (C) except for the coupled states with E/M phenotype. (E) The same as (C) except for the coupled states with E phenotype. (F) The percent of initial conditions leading to E/M, M, and E phenotypes broken down by coupled metabolic phenotype. As $\mu_{200}$ is inhibited by Hif-1, the E/M and E states are slowly suppresed until the system is nearly saturated at mesenchymal.

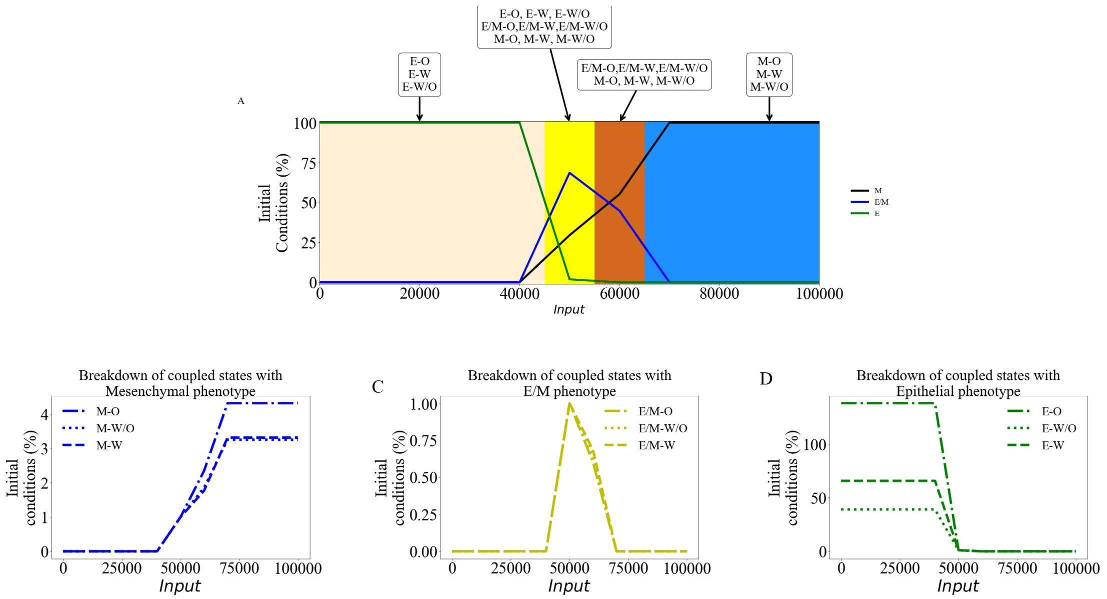


Figure 11: The input to Snail is modulated. (A) The possible sets of coupled steady states as the regulation increases. (B) The fraction of initial conditions at a particular level of regulation ($Input$) compared to the inactive state ($Input = 50000$) that lead to the M-O, M-W/O, or M-W coupled states. (C) The same as (B) except for the coupled states with E/M phenotype. (D) The same as (B) except for the coupled states with E phenotype. If the input to Snail is reduced the system saturates at fully epithelial, but if the input is increased then the system saturates at mesenchymal. Modulating the input to snail alongside other crosstalks can change the window of E/M phenotype availability.

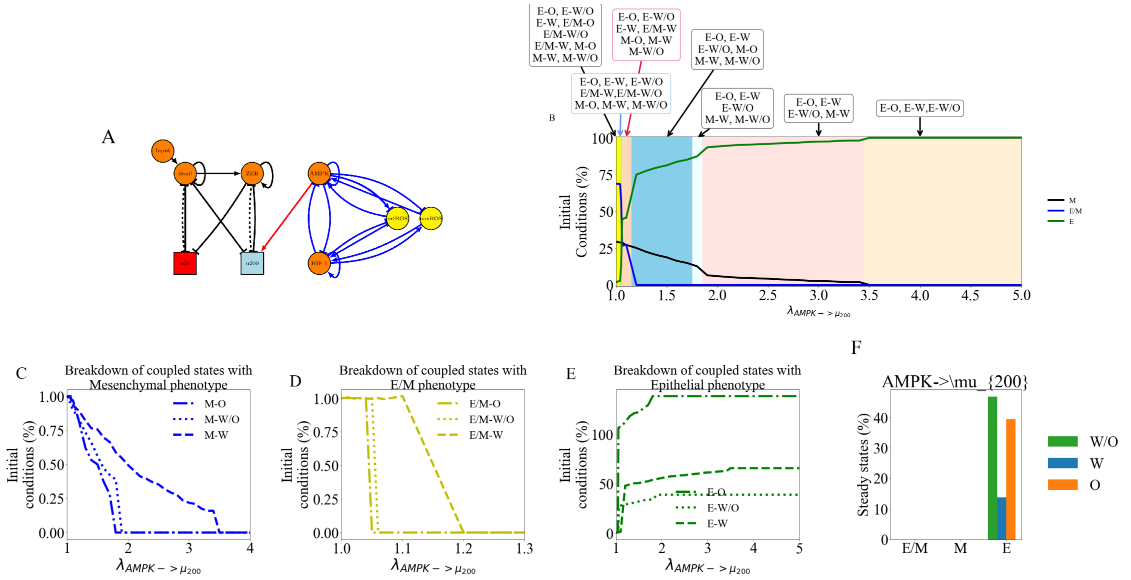


Figure 12: $AMPK->\mu_{200}$ is only active crosstalk. (A) The network showing the active crosstalk. (B) The possible sets of coupled steady states as the regulation increases. (C) The fraction of initial conditions at a particular level of regulation ($\lambda$) compared to the inactive state ($\lambda = 1$) that lead to the M-O, M-W/O, or M-W coupled states. (D) The same as (C) except for the coupled states with E/M phenotype. (E) The same as (C) except for the coupled states with E phenotype. (F) The percent of initial conditions leading to E/M, M, and E phenotypes broken down by coupled metabolic phenotype. As $\mu_{200}$ is inhibited by AMPK, the E/M and M states are slowly suppresed until the system is saturated at epithelial.

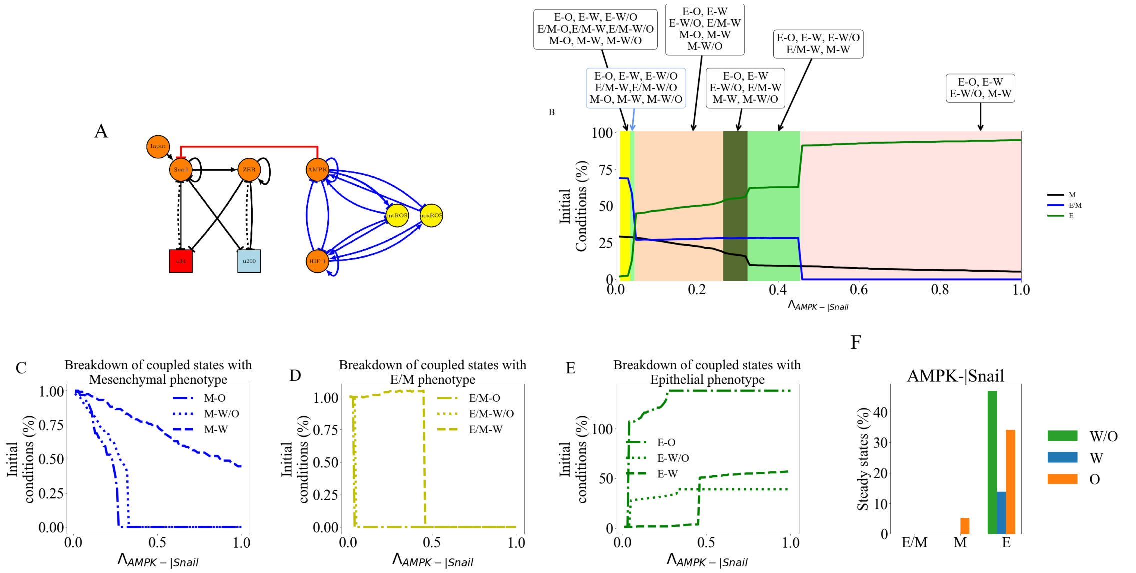


Figure 13: $AMPK - |Snail$ is only active crosstalk. (A) The network showing the active crosstalk. (B) The possible sets of coupled steady states as the regulation increases. (C) The fraction of initial conditions at a particular level of regulation ($\lambda$) compared to the inactive state ($\lambda = 1$) that lead to the M-O, M-W/O, or M-W coupled states. (D) The same as (C) except for the coupled states with E/M phenotype. (E) The same as (C) except for the coupled states with E phenotype. (F) The percent of initial conditions leading to E/M, M, and E phenotypes broken down by coupled metabolic phenotype. As Snail is inhibited by AMPK, the E/M and M states are slowly suppresed until the system is nearly saturated at epithelial.

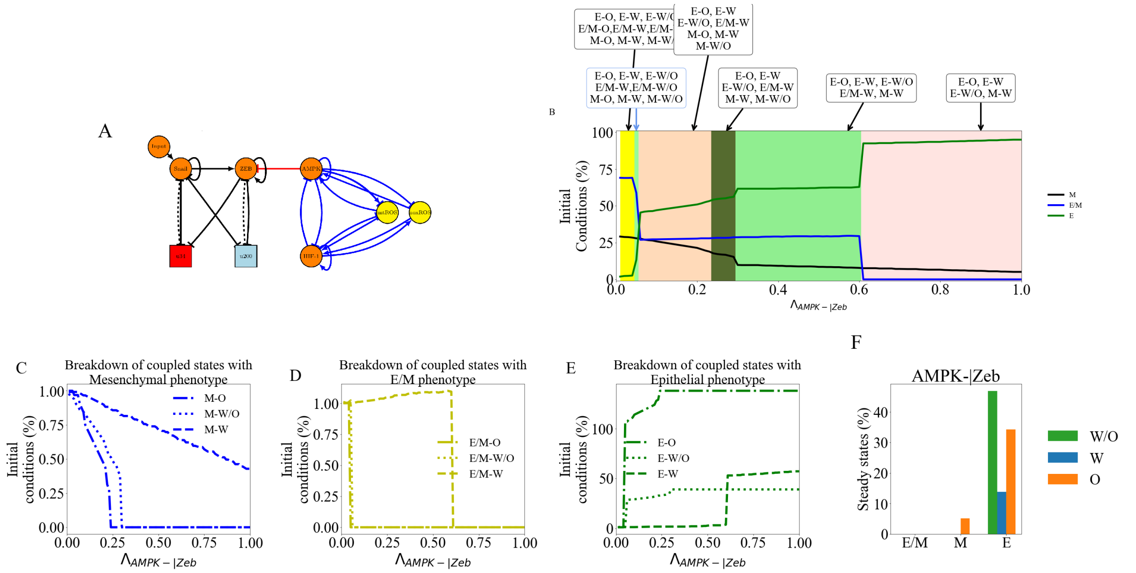


Figure 14: $AMPK - |Zeb$ is only active crosstalk. (A) The network showing the active crosstalk. (B) The possible sets of coupled steady states as the regulation increases. (C) The fraction of initial conditions at a particular level of regulation ($\lambda$) compared to the inactive state ($\lambda = 1$) that lead to the M-O, M-W/O, or M-W coupled states. (D) The same as (C) except for the coupled states with E/M phenotype. (E) The same as (C) except for the coupled states with E phenotype. (F) The percent of initial conditions leading to E/M, M, and E phenotypes broken down by coupled metabolic phenotype. As Zeb is inhibited by AMPK, the E/M and M states are slowly suppresed until the system is nearly saturated at epithelial.

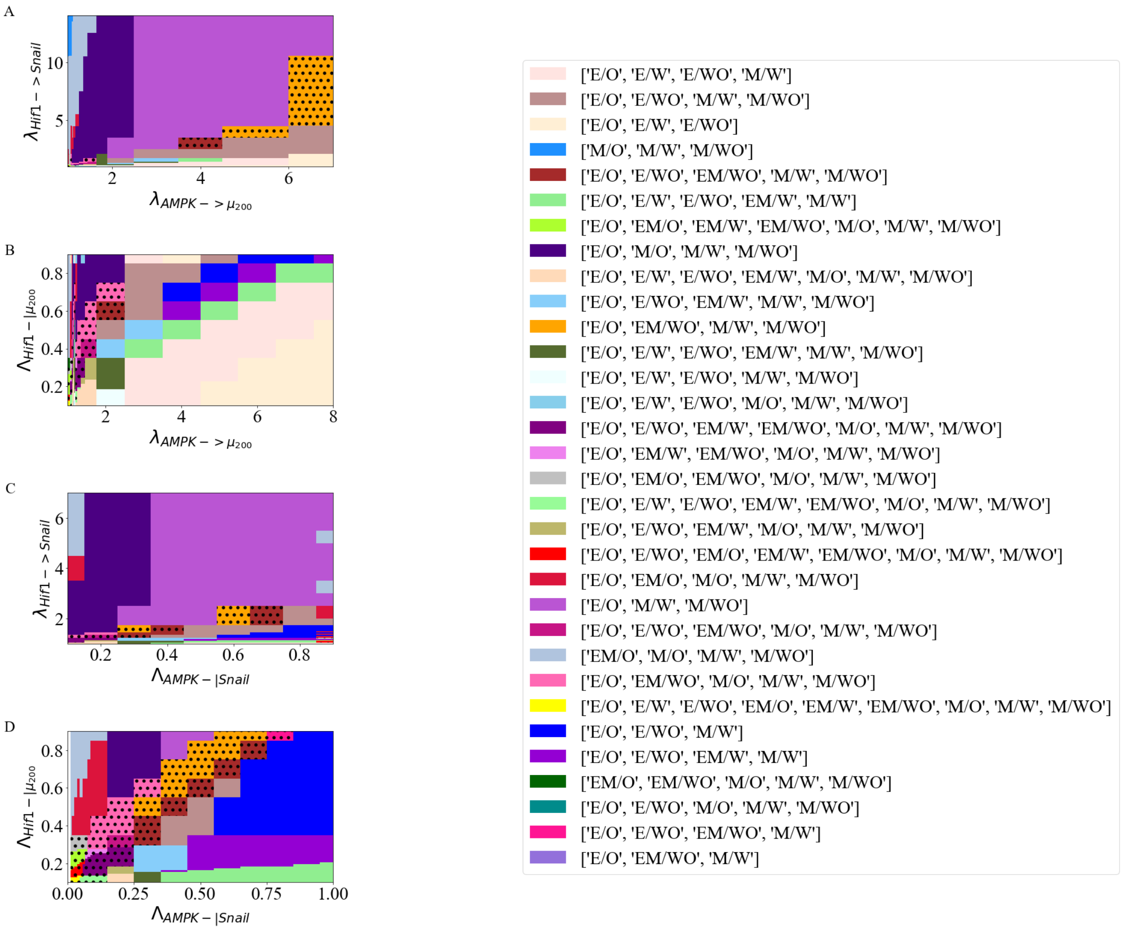


Figure 15: The potential sets of steady states when an AMPK driven crosstalk and a Hif-1 driven crosstalk are active. (A) When $Hif1->Snail$ and $AMPK->\mu_{200}$ the E/M-W/O state is more likely at high levels of regulation. (B)When $Hif1-|\mu_{200}$ and $AMPK->\mu_{200}$ the E/M-W/O state is more likely at low levels of $\lambda_{AMPK->\mu_{200}}$. (C) When $Hif1->Snail$ and $AMPK-|Snail$ the E/M-W/O state is more likely at low levels of $\lambda_{Hif1->Snail}$. (D) When $Hif1-|\mu_{200}$ and $AMPK-|Snail$ the E/M-W/O state exists when the levels of regulation by Hif1 and AMPK are similar. The results also suggest the E/M-W/O state may need to be stabilized by crosstalks acting on two different components of the network. The legend at the right is for all plots on the left.

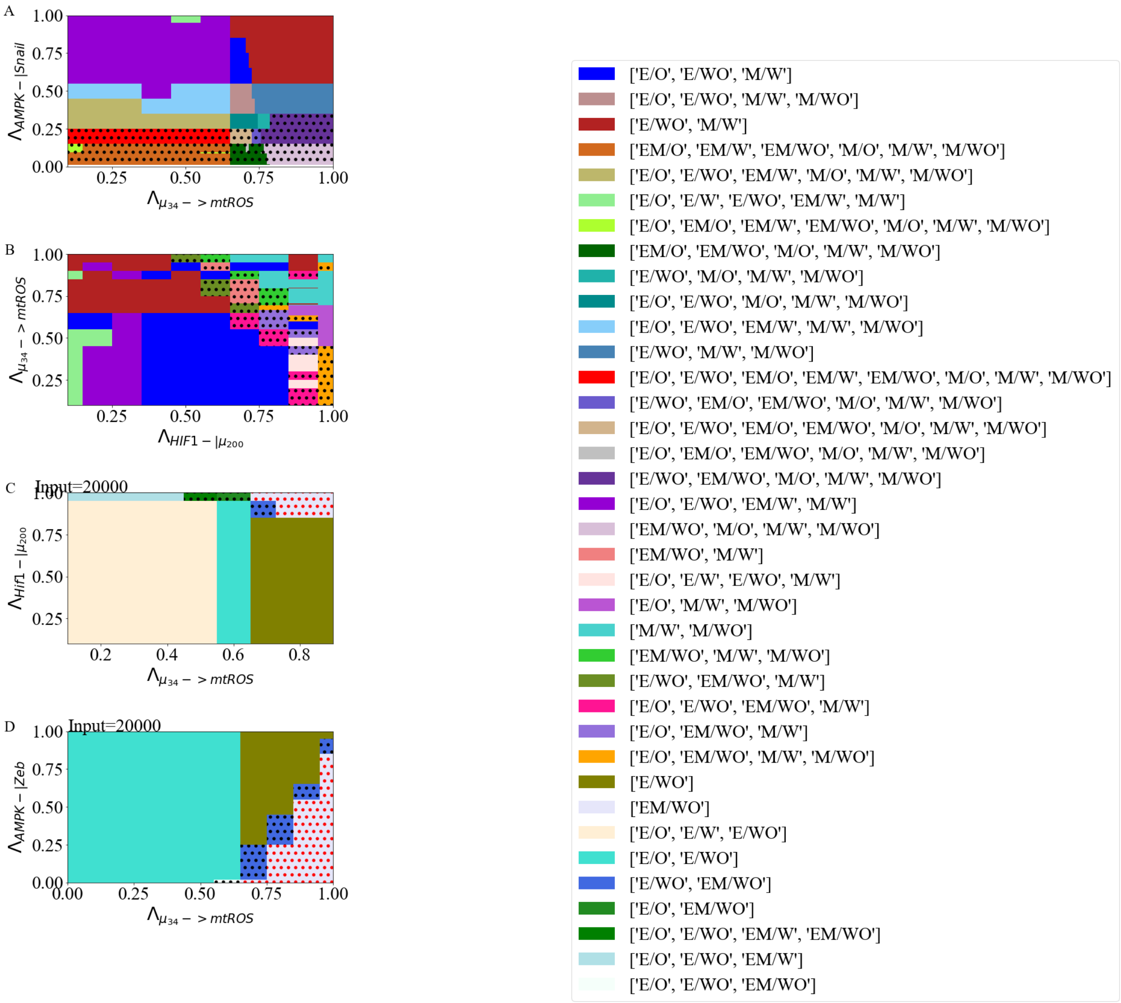


Figure 16: The figures show the entire range of parameters calculated for Fig. 6 in the main text. (A) AMPK $-|$ SNAIL, $\mu_{34}->$mtROS, and Input=20000. The E/M-W/O state is upregulated at higher levels of SNAIL. (B) AMPK $-|$ SNAIL, $\mu_{34}->$ mtROS, and HIF-1 $-|\mu_{200}$. The E/M-W/O state is upregulated when mtROS is upregulated or $\mu_{200}$ is downregulated. (C) The input to Snail is modulated, $\mu_{34}->mtROS$, and $Hif1-|\mu_{200}$ resulting in the E/M-W/O state being accessible near maximum upregulation of both mtROS and $\mu_{200}$. Additionally, the E/M-W/O state is the only one accessible at maximum regulation (near 0,0). In fact no other combination consisting of only three regulatory links seem to enable only the E/M-W/O state. (D) When all crosstalks are active (input=20000, $\lambda_{AMPK-|Snail}=0.95$, $\lambda_{AMPK->\mu_{200}}=1.1$, $\lambda_{Hif1->Snail}=1.1$, $\lambda_{Hif1-|\mu_{200}}=0.1$, UH=310).

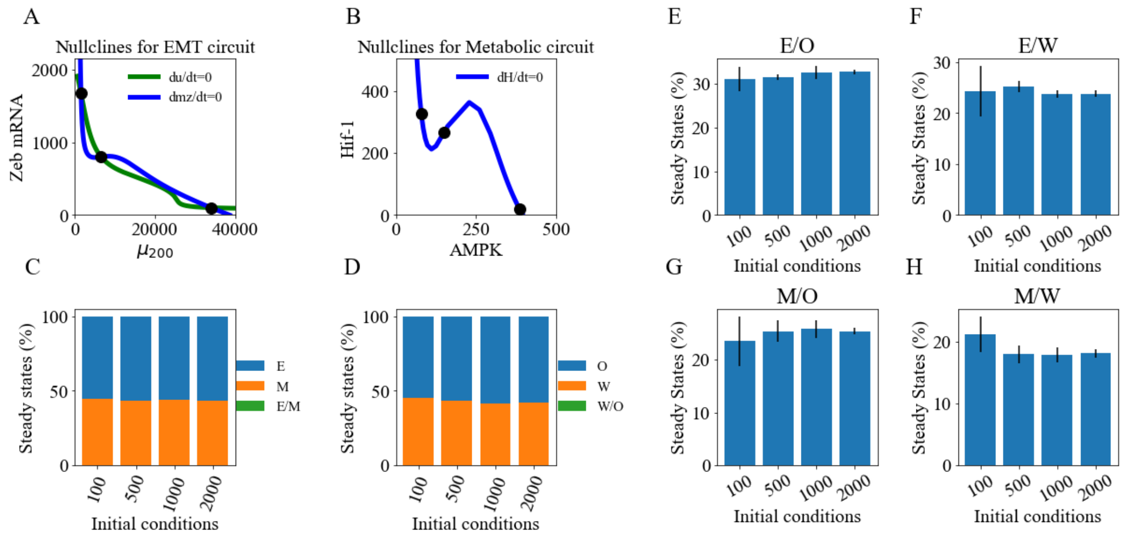


Figure 17: (A) The nullcline for the EMT network with modified parameters such that only the E and M steady states are accessible. (B) The nullcline for the metabolic network with modified parameters to ensure only the W and O steady states are accessible. (C) The percent of initial conditions leading to the E and M states (showing no E/M state) for the model with modified paramters to ensure only the E and M states are accessible. (D) The same but for the network modified to ensure no W/O state is possible. (E-H) The coupled steady states for the network modified to ensure no E/M or W/O states are initially accessible. The results show the network is almost evenly distributed across the four possible coupled states (E-O, M-O, E-W, and M-W, respectively). The results have converged once there are about 1000 initial conditions.

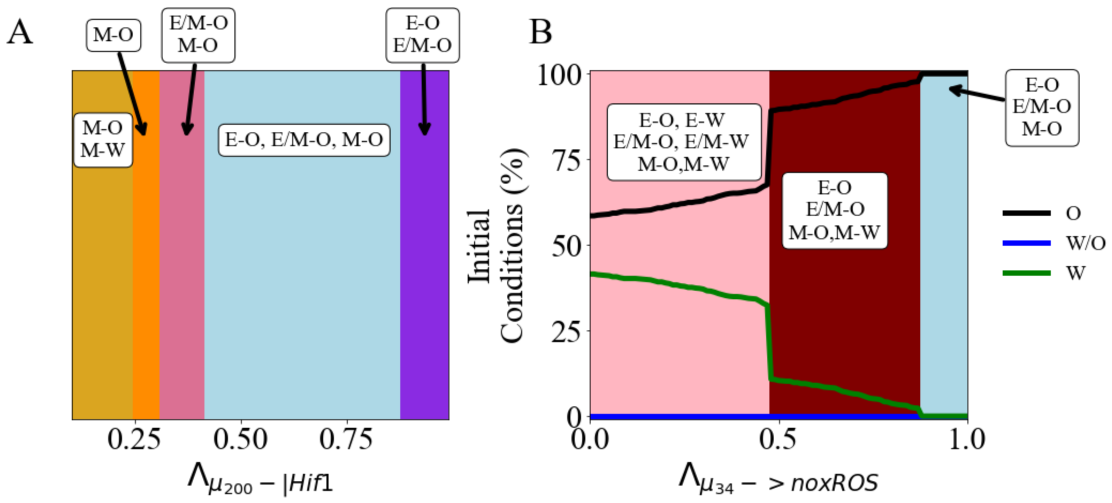


Figure 18: The network with modified parameters to ensure the hybrid W/O state is inaccessible. (A) The possible sets of steady states as the silencing of Hif1 increases, showing the system remain bistable. (B) The possibles sets of steady states as the regulation of noxROS increases showing the percent of OXPHOS metabolic states increases but the system remain bistable.

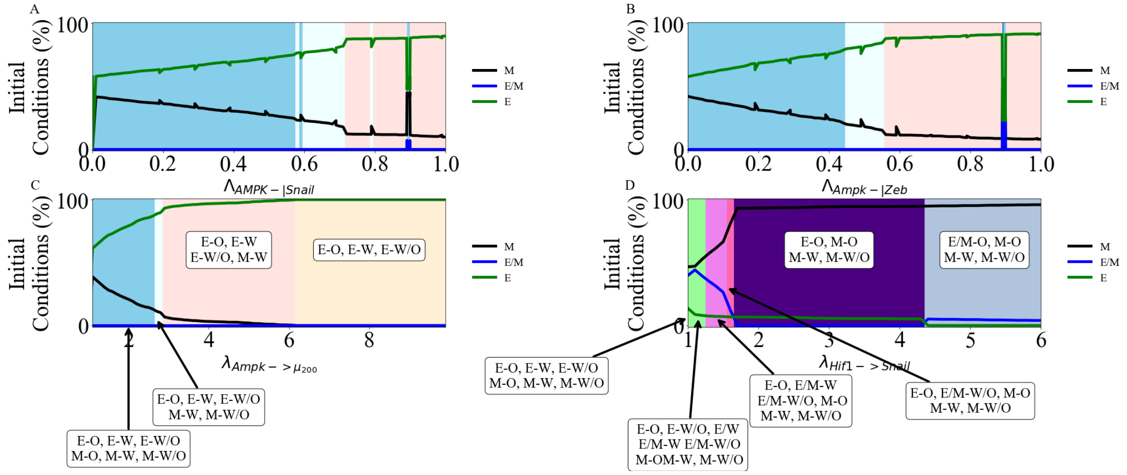


Figure 19: The network with modified parameters to ensure the hybrid E/M state is initially inaccessible. (A) As AMPK inhibits Snail, the system saturates near epithelial and remain bistable. (B) Same as A except for AMPK inhibitin Zeb. (C) Same except for AMPK upregulating $\mu_{200}$. (D) Initially, as Hif-1 slightly upregulates Snail, the E/M-W/O state is accessible but as the upregulation increases the system saturates at mesenchymal. Colors and labels are consistent across subfigures.

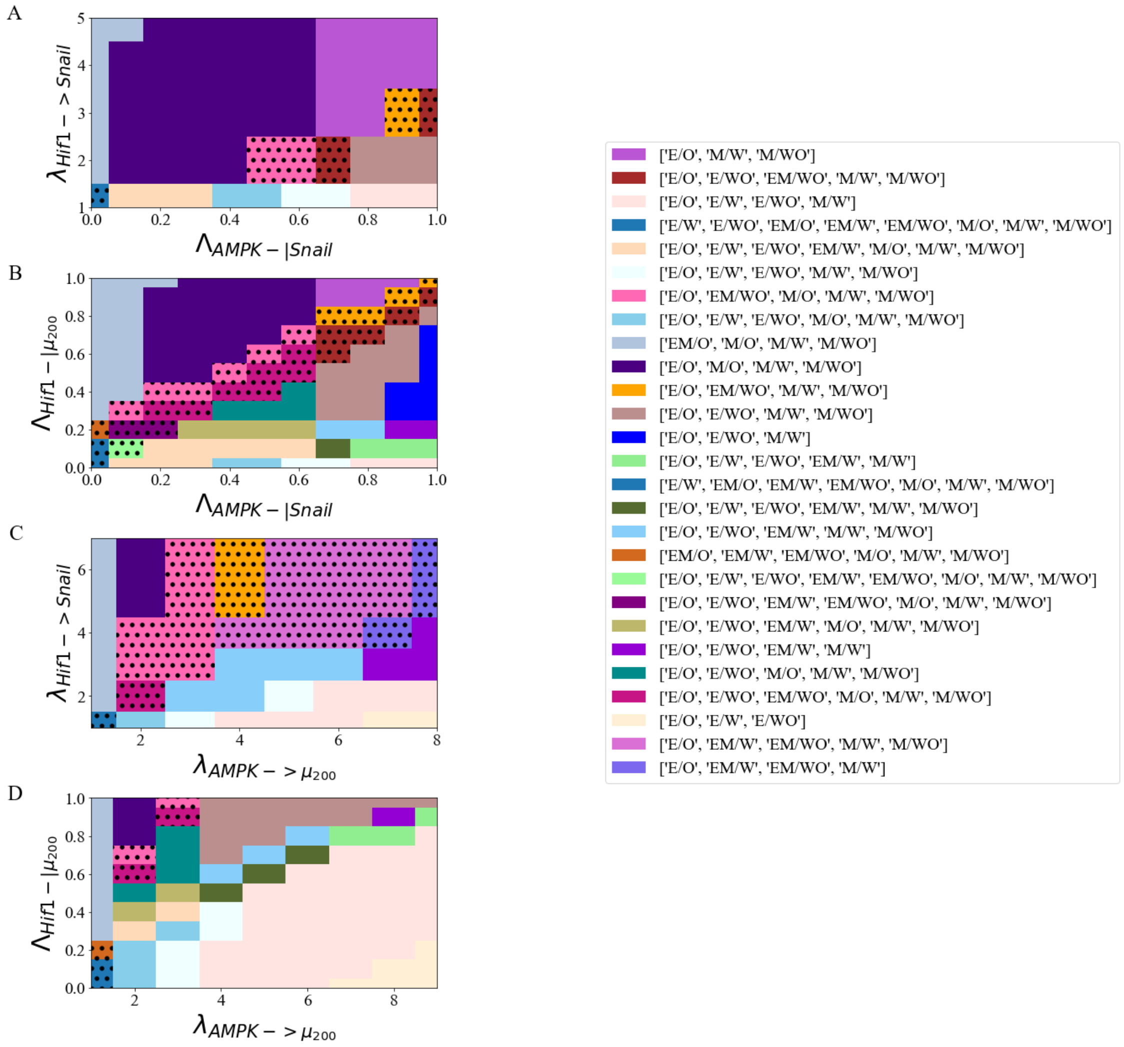


Figure 20: The network with modified parameters to ensure the hybrid E/M state is initially inaccessible. (A) When $Hif1->Snail$ and $AMPK-|Snail$ the system is able to generate the hybrid E/M state, especially when the regulation increases. (B) When $Hif1-|\mu_{200}$ and $AMPK-|Snail$ the system is able to generate the hybrid E/M state when the regulation on $\mu_{200}$ and Snail are about equal. (C) When $Hif1->Snail$ and $AMPK->\mu_{200}$ the system is able to generate the hybrid E/M state for most of the parameter ranges.(D) When $Hif1-|\mu_{200}$ and $AMPK->\mu_{200}$ the system is able to generate the hybrid E/M state, especially at low levels of $\lambda_{AMPK->\mu_{200}}$. These results show metabolic reprogramming can drive EMT.

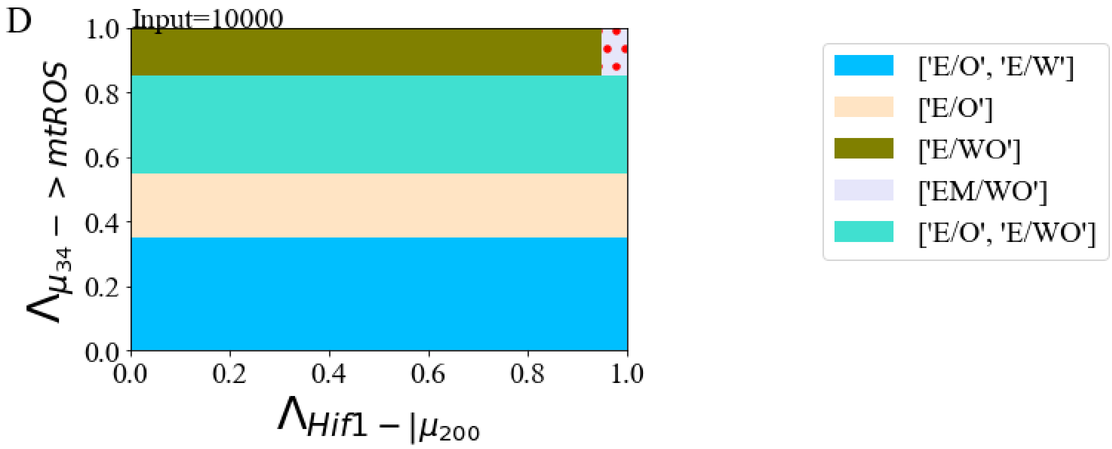


Figure 21: The full set of parameters for Fig. 8 from the main text - both networks are initially bistable (no E/M or W/O state). The E/M-W/O is able to be generated at maximum upregulationg of mtROS and inhibition of $\mu_{200}$.

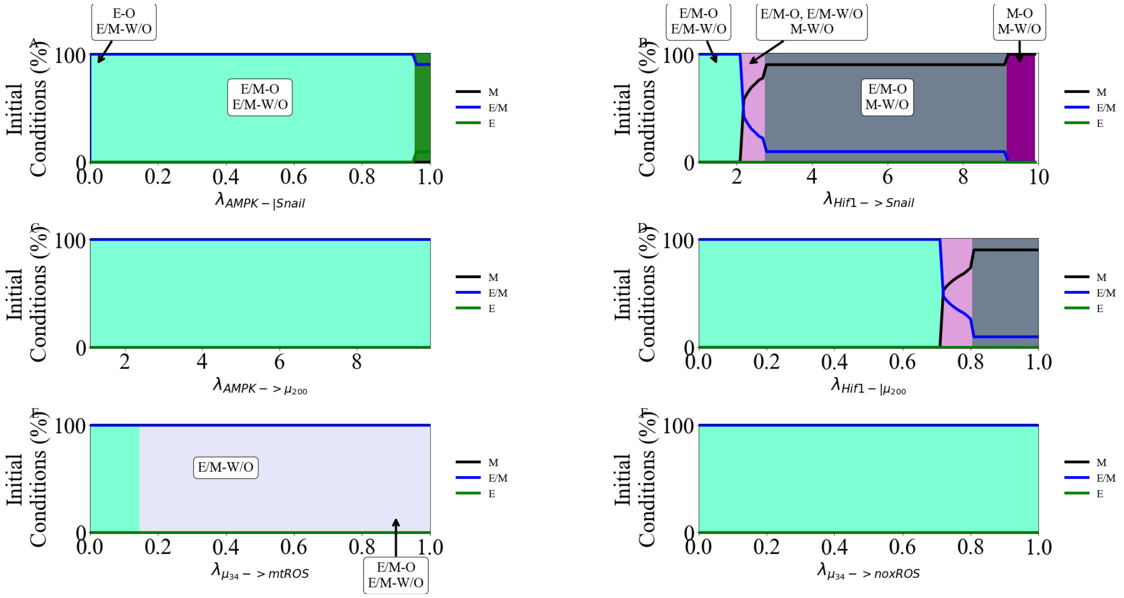


Figure 22: The activation of individual crosstalks in the network with PSFs. (A) The E/M-W/O state is maintained for all levels of $\lambda_{AMPK-|Snail}$. (B) When $Hif1- > Snail$ E/M-W/O state is accessible longer than in the tristable case before the system saturates at mesenchymal. (C) Same as (A) for $AMPK- > \mu_{200}$. (D) Same as (B) for $Hif1-|\mu_{200}$. (E) When $\mu_{34}- > mtROS$, the system saturates at E/M-W/O. (E) Same as (A) for $\mu_{34}- > noxROS$.

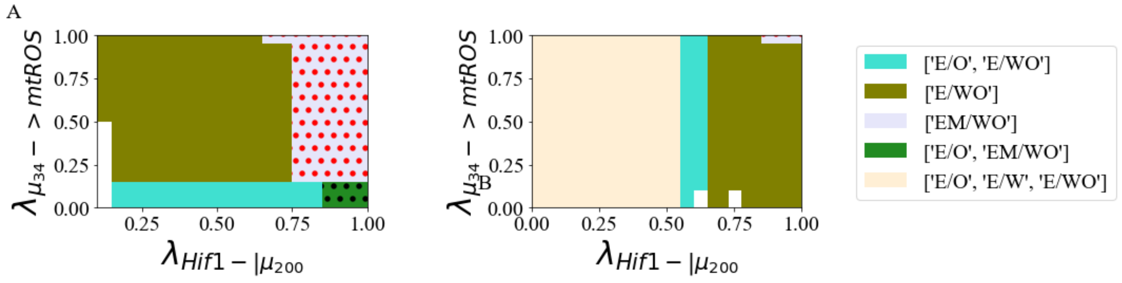


Figure 23: (A) The entire parameter range for Fig. 7c, for the network with PSFs. The input to Snail is modulated (I=10000K), $\mu_{34}->mtROS$, and $Hif1-|\mu_{200}$ resulting in approximately 20% of the parameter region only being the E/M-W/O state. (B) A zoom in near the parameter space where only the E/M-W/O state is enabled for The input to Snail is modulated (I=10000K), $\mu_{34}->mtROS$, and $Hif1-|\mu_{200}$. Both networks are initially tristable.
